# Organoids reveal niche-specific mechanotransduction-guided human cortical patterning and cell fate acquisition

**DOI:** 10.64898/2026.07.15.736390

**Authors:** Alex J. Kingston, Annabelle A. M. Wurmser, Aditi S. Kulkarni, Kejia Miao, Gökçe G. Agsu, András Lakatos, Srinjan Basu

**Author notes:** These authors contributed equally (András Lakatos, Srinjan Basu).

## Abstract

The highly dynamic mechanical environment of the developing cerebral cortex has the potential to encode information vital for neuronal differentiation. Changes in local biophysical properties have been implicated in processes ranging from the maintenance of neural progenitors to long-range axon guidance. However, whether cortical mechanics influence cellular organisation and specification during neurogenesis remains largely unexplored. Here, we leverage a 3D mosaic organoid model of human cortical development, in which we selectively disrupt a key component of nuclear mechanosensing, the LINC complex, decoupling cells from their mechanical environment. We show that LINC-decoupling alters nuclear morphology in a compartment-specific manner, driving preferential exclusion from the germinal zone and concomitant premature differentiation. Excluded cells exhibit a biased spatial distribution in the cortical plate and altered fate allocation, with loss of intermediate progenitors and upper-layer neuron populations. Combining this approach with single-cell transcriptomic profiling, we reveal a signature of impaired ERK activation and density sensing in LINC-decoupled cells. Furthermore, we show that altered density sensing contributes to mislocalisation of the nuclear envelope protein emerin and disruption of histone mark deposition during differentiation. Taken together, our findings illustrate how the dynamic mechanical environment of a complex tissue can dictate cell fate and pattern formation during development.

## INTRODUCTION

The division of labour within biological tissue drives a huge diversity of cytoarchitecture, leading to the formation of mechanically distinct regions through changes in cell packing, morphology and extracellular matrix (ECM) composition (Thompson et al. 2019, Budday et al. 2019, Singh and Chanda 2021, Ryu et al. 2021). Spatial mechanical heterogeneity is particularly apparent during development, when growth and morphogenesis exert additional bulk forces on the tissue (Heisenberg & Bellaiche 2013, Maitre et al. 2015, Chan et al. 2020). This dynamic mechanical environment encodes crucial information about the spatiotemporal context of development (Iwashita et al. 2014, Shellard and Mayor 2021, Bubna-Litic et al. 2024). However, the means by which mechanical heterogeneity of this kind affects cell type specification in a complex tissue environment remains largely unexplored.

The developing cerebral cortex contains two primary tissue architectures, a densely packed ventricular zone (VZ) and more sparsely occupied cortical plate (CP) (Figure 1A; Rakic 1988, Ramon y Cajal 1995, Lui et al. 2011). These zones possess a radial organisation which corresponds approximately to the progress of neuronal differentiation. As such, the transition from neural progenitor to neuron necessitates a concomitant change in the mechanical environment (Iwashita et al. 2014, Walter et al. 2023). Furthermore, cyclic migration of progenitors within the VZ in a process termed interkinetic nuclear migration (IKNM), and morphogenic changes associated with neuronal differentiation exert additional forces on the nucleus, transmitted via nucleocytoskeletal linkage (Reiner et al. 2011, Okamoto et al. 2013, Miyata et al. 2015, Strzyz et al. 2015). The effect of this mechanical complexity on neuronal development in human cortex is as yet unknown.

**Figure 1:**
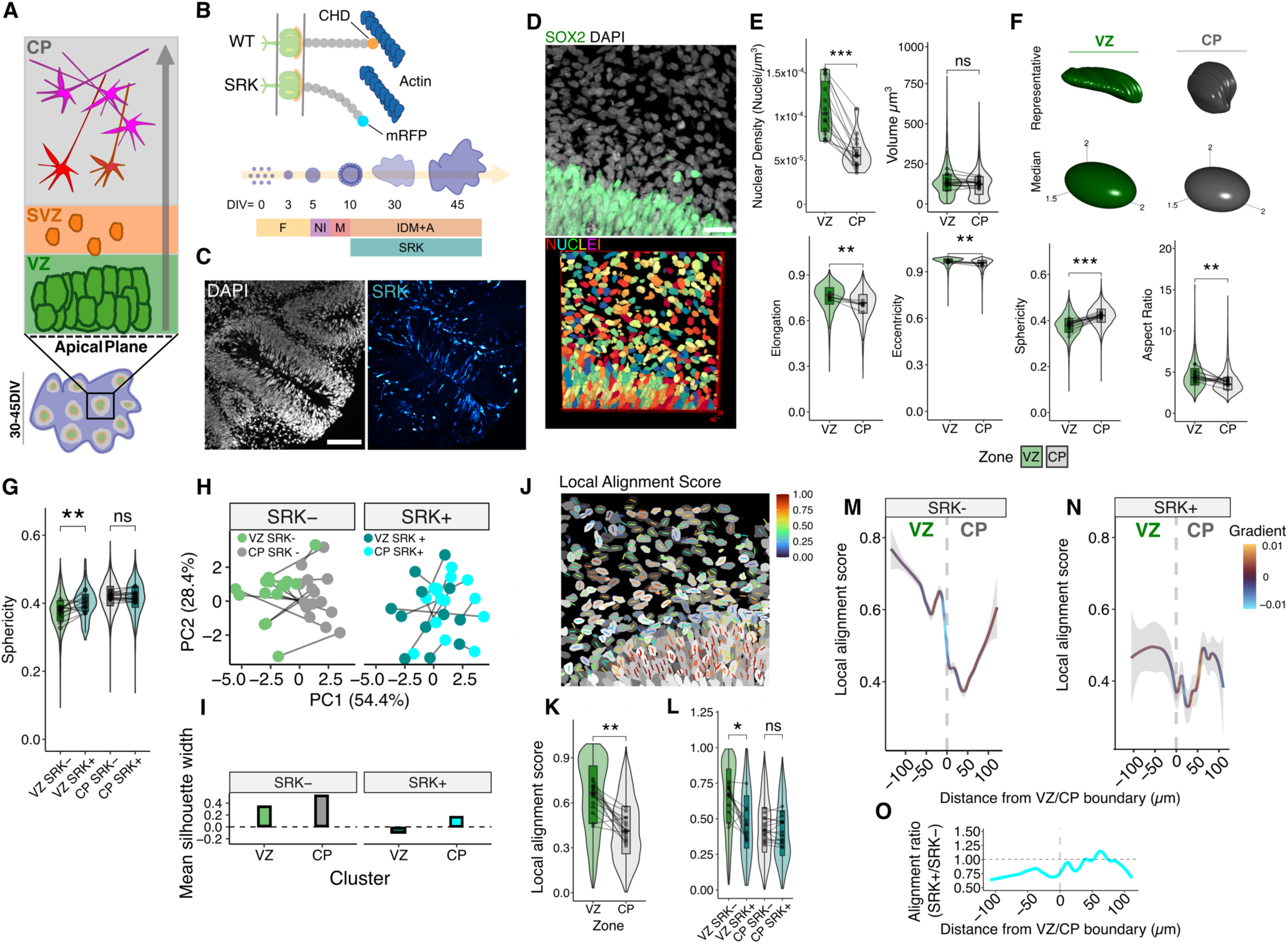
Niche-specific nuclear morphology is disrupted by LINC-decoupling. (A) Schematic of organoid tissue organisation in the ventricular zone (VZ), sub-ventricular zone (SVZ) and cortical plate (CP) along the radial axis (grey arrow). (B) Schematic of wild type (WT) and SR-KASH (SRK) constructs (top) and organoid experimental timeline (bottom). F = formation media; NI = neural induction media; M = maturation media; IDM + A = improved differentiation media + vitamin A. (C) Representative immunofluorescence (IF) images showing SRK induction in a 30DIV organoid. Scale bar 100µm. (D) Representative max-intensity projection of confocal z-stack (top) and 3D segmentation volumes (bottom) of a VZ\CP boundary. Scale bar 20µm. (E) Violin plots showing the nuclear density values, volume, and selected morphometric parameters across tissue compartments. *n* = 6 organoids (18 tissue boundaries from 3 independent differentiations). Paired two-tailed student’s t-test. From top left to bottom right *p* = 0.00051, 0.18, 0.0076, 0.0069, 0.00083, 0.0089. (F) representative (top) and median renderings (bottom) of nuclei from the VZ and CP, showing changes in gross morphology. (G) Violin plot demonstrating the sphericity values across induced cells in each tissue compartment. *n* = 6 organoids (12 tissue boundaries from 3 independent differentiations). Paired two-tailed student’s t-test. *p* = 0.0082 (VZ), 0.46 (CP) (H) PCA plot of all morphological descriptors in SRK- and SRK+ cells, showing VZ-CP transitions across all samples joined by pairing lines. *n* = 7683 cells (7012 SRK-, 683 SRK+; 12 tissue boundaries from 6 organoids and 3 independent differentiations). (I) cluster silhouette scores showing loss of cluster separation in SRK+ tissue-zone transitions. (J) Representative image of local major axis alignment, axes are plotted on corresponding nuclei and coloured by local alignment score. (K) Local alignment quantified by tissue zone in uninduced cells. *n* = 6 organoids (18 tissue boundaries from 3 independent differentiations). Paired two-tailed student’s t-test. *p* = 0.0012 (L) Local alignment relative to induced cells, showing loss of alignment in SRK+ VZ cells. *n* = 6 organoids (18 tissue boundaries from 3 independent differentiations). Paired two-tailed student’s t-test. *p* = 0.015 (VZ), 0.85 (CP) (M) Local alignment plotted as a function of VZ/CP boundary distance in uninduced cells, showing marked loss of alignment across the boundary. Ribbon depicts standard error of the mean (SEM), loess fit. *n* = 7012 cells (18 tissue boundaries spanning 6 organoids from 3 independent differentiations). (N) Equivalent plot of induced cell alignment, showing loss of VZ/CP transition. *n* = 683 cells (12 tissue boundaries spanning 6 organoids from 3 independent differentiations). (O) Ratio of mean SRK+/SRK- alignment scores over relative distance. *n* = 7683 cells (7012 SRK-, 683 SRK+; 12 SRK+/18 SRK- tissue boundaries spanning 6 organoids from 3 independent differentiations). *p*-values: * <.05, **<.01, ***<.001 or ns = not significant; Overlaid box plots show median values (line), interquartile range (box) and data range excluding outliers (whiskers).

The conversion of mechanical information into changes in cellular activity is known as mechanotransduction (Chen 2008, DuFort et al. 2011, Hoffmann et al. 2011). This process relies heavily on the nucleus, which is uniquely positioned as a mechanosensitive hub integrating cell extrinsic and intrinsic stimuli (Chalut et al. 2012, Lomakin et al. 2020, Venturini et al. 2020). One of the primary mechanisms of nuclear mechanotransduction is the direct propagation of stress waves via the Linker of Nucleoskeleton and Cytoskeleton (LINC) complex (Crisp et al. 2006, Lombardi et al. 2011, Isermann and Lammerding 2013, Alam et al. 2016). The LINC complex spans the perinuclear space and mediates physical continuity between the ECM, cytoskeleton, and intranuclear chromatin, providing a path for mechanical stress to directly alter transcription, genome topology and the deposition of epigenetic marks (Tajik et al. 2016, Nava et al. 2020, Goelzer et al 2021). Signals of this kind have previously been shown to regulate neuroectodermal competence via changes in histone modifications at the *Sox1* locus (Hamouda et al. 2025) and epidermal differentiation through transmission of integrin-dependent stress forces (Carley et al. 2021), both in 2D cell culture models. Equivalent characterisation of mechanotransduction in a complex neural tissue model is yet to be undertaken.

The advent of 3D organoid models of early tissue development has provided a means to investigate the effect of tissue-intrinsic mechanics with unprecedented accessibility and experimental tractability. These models have provided particular advantages in the study of the developing brain, given that development of the human cerebral cortex occurs largely *in utero*, typically precluding exploration of the underlying mechanical environment. Specifically, organoid models allow for controlled disruption of mechanosensation within a tissue environment (Liu et al. 2023, Weischelberger et al. 2025).

Here, we leverage cortical brain organoids (CBOs) to model the tissue architecture transitions involved in human neurogenesis (Sasai et al. 2008, Lancaster et al. 2013, Lancaster et al. 2014, Renner et al. 2017). For this, we developed cortical organoids allowing for selective LINC complex disruption. By abrogating transmission of tissue forces through, we characterise the effect of tissue-intrinsic spatial mechanical heterogeneity on neuronal commitment and cortical patterning. We find niche-specific effects on nuclear morphology indicative of altered forces acting on the nucleus, as well as accelerated timing of the transition from the VZ-CP transition and changes in the balance of mitotic division modes within the VZ. We show that LINC-decoupled cell fate and cortical positioning leads to a loss of specific progenitor and neuronal subtypes, and biased spatial distribution suggestive of disrupted differentiation, migration and cortical patterning. Single-cell transcriptional profiling of affected cells reveals signatures of altered density-sensing. Furthermore, fixed-cell cytometry reveals enrichment of lineage-specific repressive histone marks and altered localisation of the key mechanosensitive protein emerin to the outer nuclear membrane. Our results indicate that tissue-intrinsic mechanical signals relayed through the LINC complex are a key regulator of the timing and outcome of cortical neurogenesis.

## RESULTS

### Niche-specific nuclear morphology is disrupted by LINC-decoupling

To investigate how the mechanical shift from VZ to CP affects neuronal differentiation, we leveraged an inducible, red fluorescent protein (RFP)-tagged dominant-negative mutant of the LINC complex component SYNE2 (Spectrin-containing nuclear-envelope protein 2; encoding the nesprin-2 protein; Supplementary figure 1A), termed SR-KASH (SRK; Figure 1B). Doxycycline induction of SRK drives overexpression of a truncated product, lacking the calponin homology domain (CHD) needed to bind to the actin cytoskeleton (Supplementary figure 1B). SRK effectively outcompetes endogenous nesprins for binding to inner LINC components, displacing them from the nuclear membrane and abrogating force transmission to the nucleus (Luxton et al. 2010). We induced SRK expression in brain organoids at 10 days *in vitro* (DIV), after neuroectoderm differentiation, and fixed tissues after rosette formation and neurogenesis at 30-45DIV in order to examine the effect of mechanotransduction on neuronal commitment across different tissue zones using immunohistochemical labelling (Figure 1B). Sparse induction of SRK generated a mosaic tissue composition, which allowed for investigation of SRK+ cells within a predominantly wild-type environment (Figure 1C; Supplementary figure 1C).

To characterise mechanical heterogeneity within SRK-organoid tissue zones and its impact on the cell nucleus, we leveraged changes in nuclear morphology as a proxy for mechanical force (ref). We quantified 3D nuclear morphology across the VZ-CP boundary, defined by SOX2 immunoreactive (IR) cells and DAPI staining (Figure 1D; Supplementary figure 1D). As expected, we found the VZ to exhibit a far higher nuclear density than the CP (Figure 1E). Additionally, nuclei within the CP exhibited a significant increase in sphericity and concomitant decrease in aspect ratio relative to those in the VZ, without a change in overall volume (Figure 1E, 1F). We then examined the effect of SRK induction on nuclear morphology across each tissue zone, finding a VZ-specific increase in sphericity within the LINC-decoupled population (Figure 1G). Assessment of all morphological descriptors (volume, sphericity, aspect ratio, elongation, eccentricity, surface-to-volume ratio, local alignment, equivalent diameter and compactness) in PCA space revealed a drastic reduction in the separation of VZ and CP nuclei as quantified by cluster silhouette score (Figure 1H, 1I).

Next, we examined the alignment of the major axes of nuclei in different compartments (Figure 1J), prompted by recent published findings showing the effect of tissue mechanics on the organisation of neighbouring cell nuclei (Nava et al. 2020, Mcginn et al. 2021). VZ nuclei were significantly more aligned with their nearest neighbours than CP nuclei (Figure 1K), with the steepest decay in local alignment score coinciding with the VZ-CP boundary, illustrating the marked shift in mechanical environment at the tissue phase transition (Figure 1M). SRK induction drove a significant reduction in VZ local alignment (Figure 1L, characterised by a total loss of the previously observed alignment shift at the boundary (Figure 1N, 1O). Taken together, these data show that traversing the VZ-CP boundary during neuronal commitment necessitates a dramatic shift in the mechanical environment, which is transduced to the nucleus through LINC-dependent nucleocytoskeletal linkage.

### LINC-decoupling disrupts the timing of VZ delamination and cortical positioning

We next assessed if altered nuclear mechanotransduction across the VZ-CP boundary affected the timing of VZ delamination. To this end, we cloned a nuclear-envelope (NE) targeted GFP construct under an identical promoter to the SRK construct (neGFP), allowing for temporally controlled tagging of the NE without affecting mechanotransduction. We then generated chimeric organoids harbouring a mixture of SRK and neGFP cells and induced both constructs at day 10 (Figure 2C). Strikingly, we observed a higher proportion of SRK+ cells excluded at earlier timepoints (Figure 2D), suggesting VZ retention and self-renewal is compromised in induced cells (Figure 2E).

**Figure 2:**
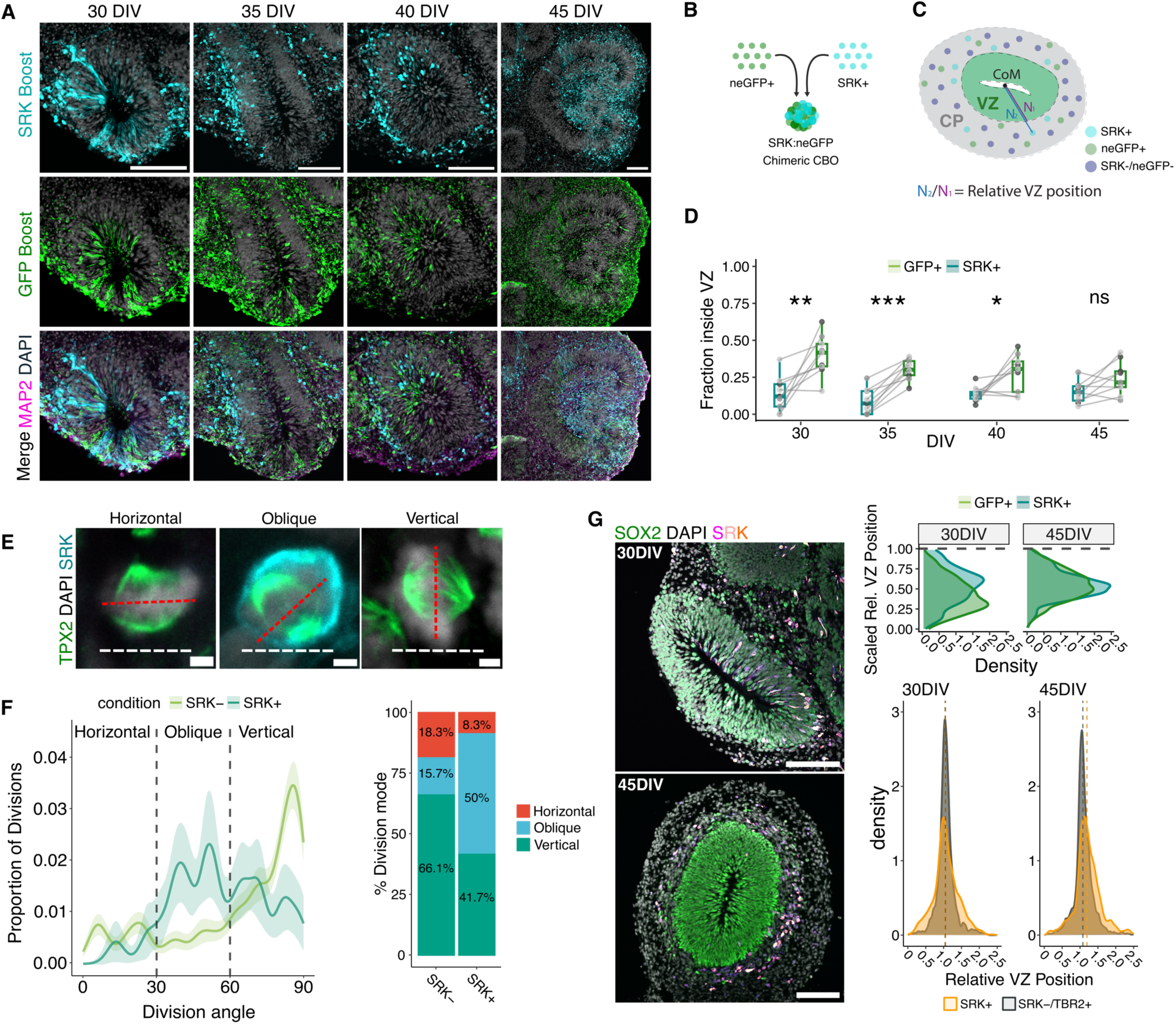
LINC-decoupling disrupts the timing of VZ delamination and cortical positioning. (A) Representative images of chimeric VZs at 30, 35, 40 and 45DIV. Scale bars 100µm. (B) Schematic of chimeric organoid generation to track VZ exclusion rate. (C) Schematic depicting quantification of relative VZ position, and cortical position (scaled by maximum recorded N2 per organoid) used in positional quantifications. (D) Quantification of VZ-retained proportion in SRK+ vs neGFP+ cells at each timepoint. *n* = 9 tissue boundaries per timepoint from 3 organoids. Paired two-tailed student’s t-test. From left to right *p* = 0.0054, 0.000057, 0.026, 0.054. (E) Representative images showing mitotic division angles relative to the apical VZ surface. Scale bar 2µm. (F) Plot depicting distribution of division angles in SRK+ and SRK- radial glia (left) and percentages of horizontal, vertical and oblique divisions (right). Ribbon depicts standard error of the mean, loess fit. *n* = 115 SRK-/24 SRK+ cells (329 VZs from 7 organoids and 3 independent differentiations). (G) Representative image of SRK+ cell accumulation in the SVZ at 30 and 45DIV (left, scale bars 100µm), density plots depicting accumulation of SRK+ cells within the SVZ, and VZ position overlap of SRK+ population with endogenous TBR2+ intermediate progenitors at 30 and 45DIV. Upper panel 30 DIV: *n* = 840/889 GFP+/ SRK+ cells 9 VZs from 3 organoids). 45 DIV: *n* = 2309/3327 GFP+/ SRK+ cells (9 VZs from 3 organoids). Lower panel 30DIV: n = 1197 TBR2+/712 SRK+ cells (8 control/8 induced organoids and 32 control/59 induced VZs from 3 independent differentiations). 45 DIV: *n* = 4179 TBR2+/2246 SRK+ cells (8 control/8 induced organoids and 36 control/99 induced VZs from 3 independent differentiations). *p*-values: * <.05, **<.01, ***<.001 or ns = not significant; Box plots show median values (line), interquartile range (box) and data range excluding outliers (whiskers).

The orientation of mitotic cleavage relative to the apical plane of the VZ correlates with the timing of VZ delamination and neuronal differentiation, as well as relying on tissue scale mechanical force to regulate its orientation (Lamonica et al. 2013, Lancaster et al. 2013, Lam et al. 2020). To investigate if the observed premature delamination was reflected at this level, we analysed division angles across SRK+ and SRK- radial glia at 30DIV (Figure 2E). We found a marked bias towards oblique and horizontal divisions in SRK+ cells, associated with premature neurogenesis and delamination from the VZ (Figure 2F). Given these effects on VZ retention, we assessed SRK+ cell positioning at later timepoints and observed a marked accumulation of cells at the VZ border emerging between 30 and 45DIV, in a region analogous to the *in-vivo* sub-ventricular zone (SVZ; Figure 2G). Quantification of SRK+ cell-positioning relative to TBR2-positive endogenous intermediate progenitors, canonically present in the SVZ, confirmed this overlap (Figure 2G; Supplementary figure 2A). These data support a role for nuclear mechanotransduction in the early coordination of spatial organisation and neurogenic timing during early cortical development.

### LINC-decoupled cells are not competent to produce IPs and display biased neurogenic potential

To investigate the effect of SRK induction on cell fate, we performed immunohistochemical analysis of SRK+ cells in the SVZ at 45DIV (Figure 3). Despite their spatial colocalisation with TBR2+ IPs, TBR2 IR was significantly depleted in SRK+ cells (Figure 3A; Supplementary figure 2A). This loss of IP fate was also apparent in SRK+ cells at 15 DIV, the earliest onset of TBR2 expression in the tissue, suggesting that LINC-decoupling precludes IP fate acquisition rather than impacting fate maintenance (Supplementary figure 2B). Interrogation of further cell-fate markers showed that the majority of SRK+ cells at 45DIV had acquired a neuronal fate, as evidenced by expression of the canonical marker of neuronal commitment TBR1 (Figure 3B). However, neuronal subtype-specific analysis revealed a bias in the acquisition of neuronal fates, whereby the SRK+ population was overwhelmingly positive for markers of early-born deep-layer neurons (TBR1, CTIP2), and conversely depleted for markers of later-born, more superficial layers (SATB1) relative to uninduced cells in the CP (Figure 3B). To establish if this effect was due to a developmental delay in upper layer neurogenesis within the SRK+ population, we examined samples at 60DIV, finding no recovery in upper-layer neuron specification, while deep-layer neuron specification remained unaffected (Supplementary figure 2C). These data suggest LINC-mediated nuclear mechanotransduction is a crucial regulator of progenitor fate in the developing cortex.

**Figure 3:**
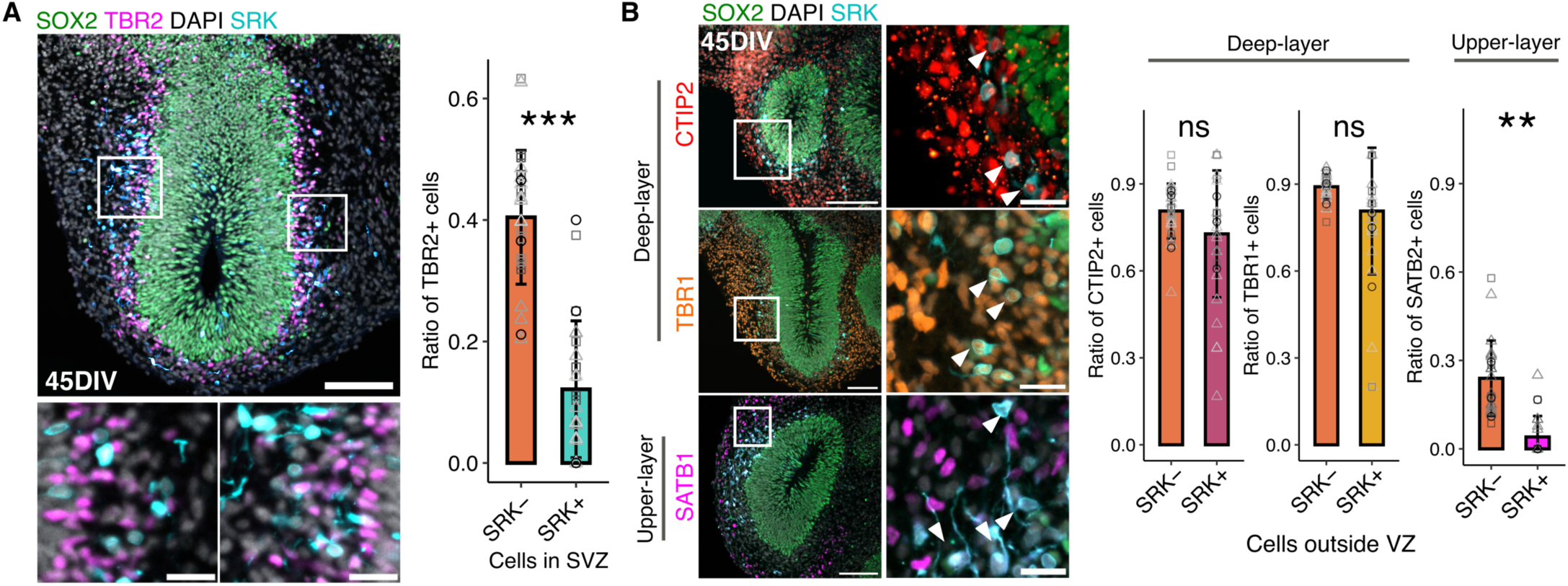
LINC-decoupled cells are not competent to produce IPs and display biased neurogenic potential. (A) Representative image at 45DIV showing SRK and TBR2 expression (left), and quantification of TBR2 expression in the SVZ between SRK+ and SRK- cells (right). *n* = 8 organoids (26 CPs from 3 independent differentiations). Wilcoxon test. *p* = 0.00094. Scale bars 100µm (overview) 50µm (inset). (B) Representative images of TBR1, CTIP2 and SATB1 expression, co-stained for SRK (left), quantifications of expression of each marker in the CP between SRK+ and SRK- cells (right). *n* = 7 organoids (33 (TBR1), 29 (CTIP2), 20 (SATB2) CPs from 3 independent differentiations). Wilcoxon test. *p* = 0.2 (TBR1), 0.066 (CTIP2), 0.0049 (SATB2). *p*-values: **<.01, ***<.001 or ns = not significant; Bars show mean values and error bars show ± standard deviation. Scale bars 100µm (overview) 50µm (inset).

### scRNA-sequencing analysis reveals altered differentiation trajectories and ERK crowding response in LINC-decoupled cells

To uncover the molecular mechanisms by which LINC decoupling affects neurogenesis, we performed single-cell RNA-sequencing (scRNA-seq) of organoid tissues at timepoints spanning the onset of upper-layer neurogenesis (Figure 4A). We profiled 10,006 cells from induced and uninduced tissues at 30 and 45 DIV using plate-based combinatorial split-pool barcoding (Figure 4B; Rosenberg et al. 2018). Unsupervised clustering and label-transfer-based annotation of clusters using an *in-vivo* human fetal brain atlas yielded 7 main cell-type clusters (Figure 4A, Supplementary figure 3A, 3B, 3C). As expected, 45DIV samples yielded higher proportions of differentiated cell types and depletion in progenitor clusters relative to 30 DIV (Supplementary figure 3D). Conversely, comparison of cluster proportions between treatment groups at both timepoints yielded no differences when assessed with differential abundance testing, likely due to the low proportion of SRK induction within the tissue (Supplementary figure 3D, Dann et al. 2022).

**Figure 4:**
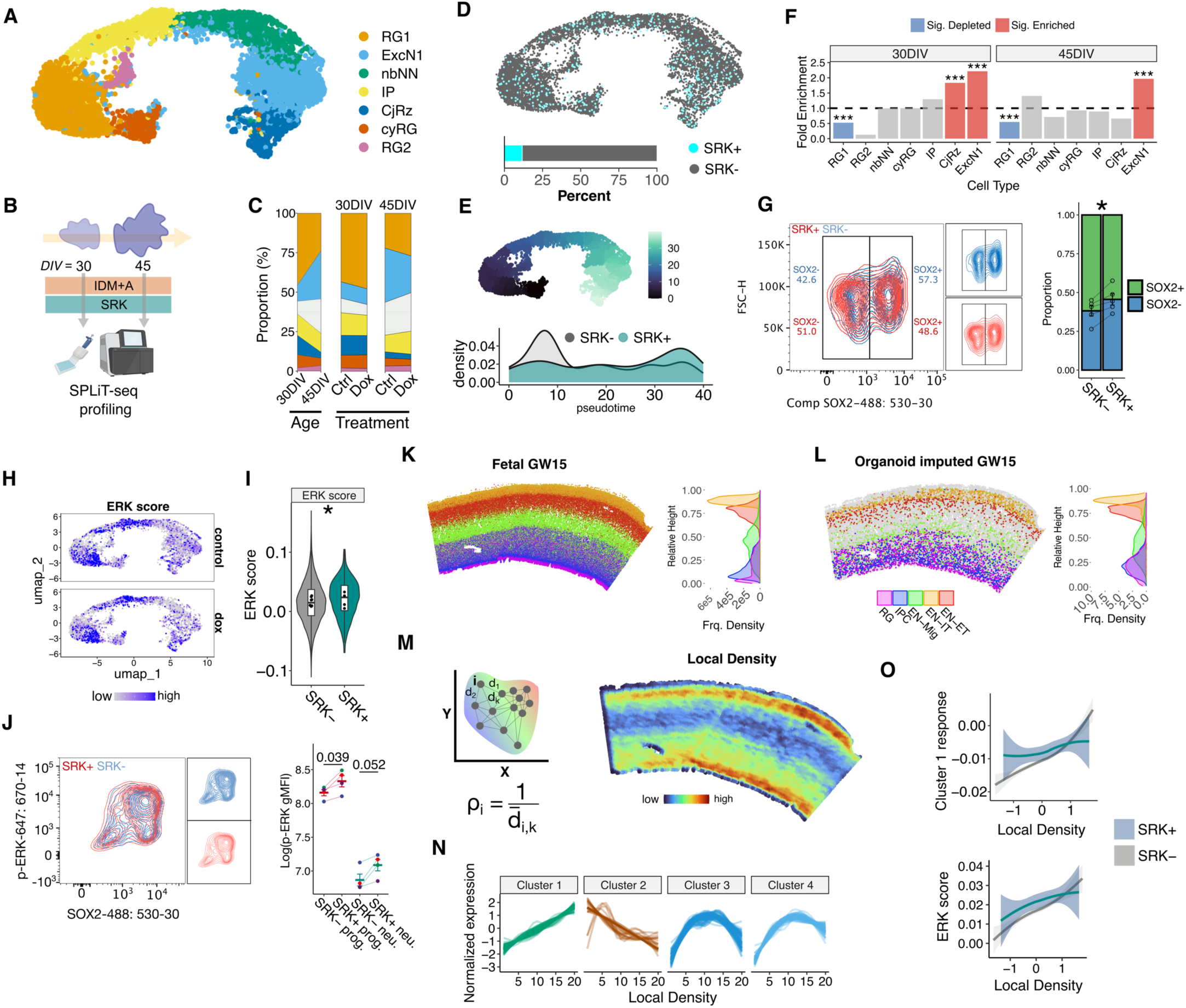
scRNAseq reveals altered differentiation trajectories and ERK-mediated crowding response in LINC-decoupled cells. (A) CCA-integrated UMAP embedding of organoid scRNAseq data coloured by cell type cluster. (B) Schematic of sample collection for scRNAseq. (C) Proportion of cell types in each sample split by timepoint (left) and treatment group (right). (D) UMAP projection of dox-treated samples highlighting SRK+ cells. (E) Density ridgeplot showing distribution of SRK+ and SRK- cells along pseudotime. (F) Fold enrichment of SRK+ cells in each cell type, per timepoint. *N* = 4 samples from 2 independent differentiations. Fisher’s exact test. From left to right *p* = 1.76e-10, 0.00056, 0.45, 0.99, 0.99, 4.48e-13, 0.42, 4.48e-13, 0.42, 2.38e-12, 0.54, 1, 0.088, 0.088, 0.52, 7.87e-06. (G) Stacked barchart showing enrichment of SOX2- progeny in SRK+ population. Samples from *n* = 5 independent differentiations. Paired two-tailed student’s t-test. *p* = 0.011. (H) Composite expression of known ERK targets across treatment groups, quantified in (I). (I) *n* = 4 samples from 2 independent differentiations. Linear model with age as covariate, *p* = 0.036. (J) Fixed-cell cytometry data showing an increase in mean p-ERK signal in SRK+ cells and quantification. Samples from *n* = 4 independent differentiations. Paired two-tailed student’s t-test. *p* = 0.039 (progenitors), 0.051 (neurons). (K) Representative spatial embedding of gw15 merFISH data coloured by cell type (right), ridgeplot showing relative spatial distribution of each cell type (left). (L) Organoid cells projected onto fetal data by imputed spatial coordinates, coloured by inferred atlas cell type (right), ridgeplot showing relative spatial distribution of each celltype (left). (M) Schematic of local density calculation from merFISH cell coordinates (left), local density map of gw15 fetal data (right). (N) Module scores plotted against local density for tradeSeq genes clustered by similarity of response pattern. Fit with a generalised additive model. (O) Lineplots depicting relationship between imputed local density and mean cluster 1 gene expression (top) or ERK target expression (bottom) in SRK- and SRK+ cells. Ribbon depicts standard error of the mean, loess fit. *p*-values: * <.05, **<.01, ***<.001; Overlaid box plots show median values (line), interquartile range (box) and data range excluding outliers (whiskers).

We therefore directly identified SRK-induced cells within our dataset by mapping reads to a custom reference genome containing the exogenous portion of the SRK construct sequence, annotating cells with detectable reads in dox-treated samples as SRK+ (Figure 4D). Approximately 10% of dox-treated cells expressed an SRK transcript, which we verified by FACS for RFP fluorescence, showing consistent cell proportions (Supplementary figure 3E, 4F). Trajectory analysis using monocle3 showed an accumulation of SRK+ cells at later pseudotime values relative to uninduced cells (Figure 4E; Trapnell et al. 2014), which corresponded to significant enrichment in neuronal clusters and depletion in the largest radial glial cluster at both timepoints (Figure 4F). For further validation, we used fixed-cell cytometry that confirmed a significant depletion in SOX2+ cells in the SRK-expressing population at 30DIV (Figure 4G). Altogether, our data suggest that LINC-decoupled cells are subject to premature differentiation when excluded from the VZ.

To explore the potential functional consequences of LINC decoupling, we carried out weighted gene co-expression network analysis (WGCNA; Morabito et al. 2023; Supplementary figure 3G). WGCNA identified 7 total gene modules, one of which, module 2, was significantly downregulated in dox-treated samples relative to control (Figure 4H; Supplementary figure 3H, 3I). Expression of this module correlated with the onset of neurogenesis when scored across the UMAP projection, and top contributors to the module eigengene included master neurogenic regulators and markers of neuronal maturity (Supplementary figure 3J). Gene set enrichment analysis of all component genes confirmed this association, with top ontology terms identified spanning synaptic maturation, neuronal morphogenesis and projection guidance (Supplementary figure 3K). These data suggest that LINC-decoupling impairs neuronal maturation processes, consistent with altered neurogenesis in the SRK+ population.

Based on the observed cell fate changes, we hypothesised that LINC-decoupling would drive detectable changes in the signature of canonical mechanosensitive transcription factors (TFs). Foremost amongst such factors are Yes-associated protein (YAP) and Extracellular-signal related kinase (ERK), both of which have been shown to regulate neurogenesis and alter their localisation and transcriptional activity in response to changes in environmental stiffness and local crowding (Panciera et al. 2017, Crozet and Levayer 2023, Jain et al. 2025). We assembled a list of known targets of each TF and generated compound scores of target expression within our dataset (Cordenonsi et al. 2011, Chen et al. 2023). We found no detectable change in expression of canonical YAP targets (Supplementary figure 3L). Conversely, ERK target expression was significantly increased across all SRK induced cells, but not between conditions alone (Figure H, 4I, Supplementary figure 3M). To validate this, we profiled ERK phosphorylation in SRK+ cells using fixed cell cytometry, through which we found a significant phosphorylation increase indicative of hyperactivation (Figure 4J).

Next, we set out to determine if this aberrant TF activity might be linked to local mechanics of the developing human brain. To this end, we utilised a published mosaic-integration approach to integrate our dataset with a merFISH atlas of human pre-frontal cortex (PFC) development spanning gestational weeks 15-22 (Ghazanfar et al. 2021, Harland et al. 2024, Qian et al. 2025; Supplementary figure 4A). After projecting organoid and reference cells onto a shared latent space, we imputed spatial coordinates for organoid cells based on their nearest neighbours in each reference sample (Figure 4K, 4L; Supplementary figure 4B). We subsequently calculated a local density score within each sample by averaging the Euclidean distance between each cell centroid and its 10 nearest neighbours (Figure 4M; Supplementary figure 4B). We imputed local density scores across organoid cells and extracted density-responsive genes using the tradeseq package (Van den Berge et al. 2020; Supplementary figure 4C). By clustering genes with similar response patterns, we identified a large cluster of responsive genes with expression proportional to local cell density (Cluster 1, Figure 4N; Supplementary figure 4D). Strikingly, this relationship was disrupted in SRK+ cells, which displayed tonically elevated expression of cluster 1 genes regardless of local density (Figure 1O). Furthermore, we observed that the previously calculated ERK response score displayed a similar relationship - with SRK+ cells displaying a reduced response to imputed local density, despite the two scores sharing <1% of component genes (Figure 1O; Supplementary figure 4E). Taken together, these data show that disrupting the sensing of local tissue mechanics impacts cortical cell differentiation and maturation through the altered activation of canonical mechanosensitive TFs.

### LINC decoupling drives aberrant transcription and widespread epigenomic dysregulation via emerin mislocalisation

We noted that differential expression analysis of our scRNAseq clusters between induced and uninduced cells yielded a neuron-specific upregulation of transcription (Figure 5A), consistent with previous work showing mechanical regulation of bulk transcription (Le et al. 2016). We corroborated this observation by fixed-cell cytometry, gating cells for SOX2 expression and finding an enrichment in RNA polymerase II phosphorylation at the serine-2 residue in SRK+/SOX2- cells, indicative of increased transcriptional activation in the non-progenitor population relative to uninduced counterparts (Figure 5B).

**Figure 5:**
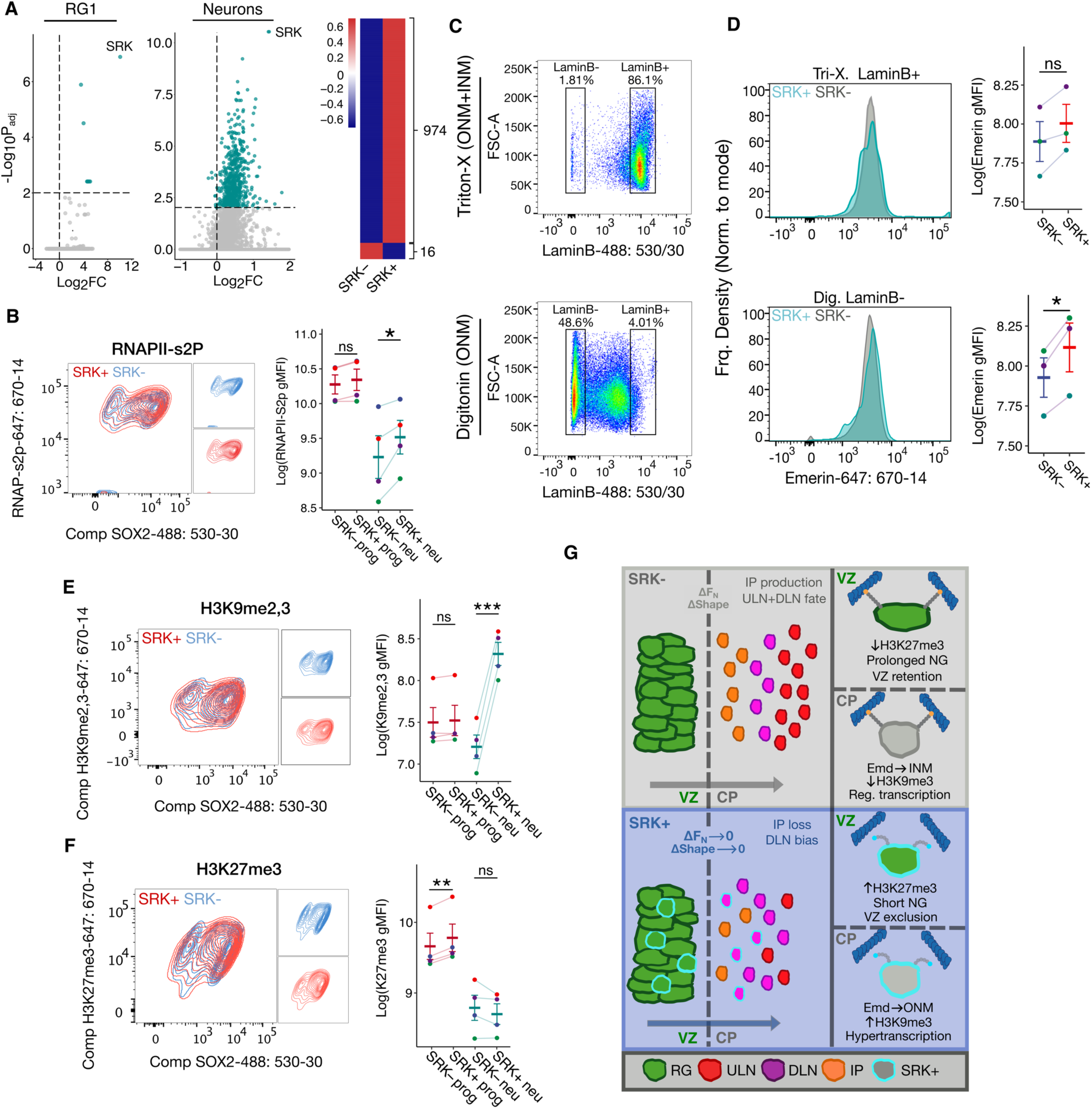
LINC decoupling drives widespread epigenomic dysregulation via Emerin mislocalisation. (A) DEGs in SRK+ vs SRK- radial glia and neurons from dox-treated scRNAseq data (left) and heatmap showing all significant DEGs from neuronal clusters (*p.adj* < 0.05), highlighting global transcriptional upregulation (right). Wilcoxon rank-sum test. (B) Fixed-cell cytometry plots of RNAPII serine-2 phosphorylation in SRK+ and SRK- populations (left) and quantification (right). Samples from *n* = 4 independent differentiations. Paired two-tailed student’s t-test. *p* = 0.055 (progenitors), 0.047 (neurons). (C) Representative scatter plots of cytometry data showing gating for LaminB+ and LaminB-populations in each permeabilisation condition. (D) Density plots showing Emerin signal in SRK+ and SRK- populations in each permeabilisation condition and corresponding quantification. Samples from *n* = 3 independent differentiations. Paired two-tailed student’s t-test. *p* = 0.076 (Triton-X), 0.028 (Digitonin). (E) Fixed-cell cytometry data of H3K27me3 and (F) H3K9me2,3, showing celltype-specific alterations in the deposition of methylation marks. Samples from *n* = 4 independent differentiations. Paired two-tailed student’s t-test. H3K9me2,3: *p* = 0.13 (progenitors), 0.0001 (neurons). H3K27me3: *p* = 0.0012 (progenitors), 0.17 (neurons). (G) Model schematic. *p*-values: * <.05, **<.01, ***<.001 or ns = not significant; Overlaid box plots show median values (line), interquartile range (box) and data range excluding outliers (whiskers).

We next investigated the localisation of the nuclear envelope protein emerin, which switches between outer and inner nuclear membranes (ONM/INM) in response to extrinsic forces and has been mechanistically linked to similar changes in bulk transcription (Le et al. 2016, Towbin et al. 2013, Morano and Holaska 2022). We observed that emerin IR at the NE increases during the VZ-CP progression despite minimal change in transcript abundance over pseudotime, suggesting recruitment of the complex to the NE correlates with neuronal commitment (Supplementary figure 5B, 5C).

To assess INM vs ONM localisation, we employed an established selective permeabilisation approach (Adam et al. 1990), whereby cells were treated with either a strong (0.1% Triton-X) or weak (0.001% Digitonin) detergent, allowing staining of both nuclear membranes or the ONM only, respectively (Supplementary figure 5A). By gating for Lamin B signal, which is uniformly expressed and localised exclusively at the INM, we were able to identify a partially permeabilised cell population in the digitonin-treated condition (Figure 5C). SRK+ cells showed no change in overall emerin signal relative to uninduced cells, but a significant enrichment in ONM-specific signal, implying a switch in emerin membrane localisation driven by LINC-decoupling in the tissue (Figure 5D).

We subsequently profiled the deposition of two histone marks known to be regulated by emerin relocalisation, H3K9me2,3 and H3K27me3 - which mark constitutive and facultative heterochromatin, respectively (Demmerle et al. 2012, Le et al. 2016, Hamouda et. al 2025). Consistent with previous studies, we observed effects on deposition of both marks in LINC-decoupled cells, with a markedly stronger effect on H3K9me2,3 deposition (Demmerle et al. 2012; Figure 5E, 5F; Supplementary figure 5D). Examining deposition patterns within neuronal and progenitor populations revealed a population-specific pattern of enrichment. SOX2+ progenitors displayed no change in H3K9me2,3 signal, combined with a modest enrichment in H3K27me3. Conversely, the SOX2- population showed a drastic enrichment in H3K9me2,3 with no change in H3K27me3 (Figure 5E, 5F). In summary, our data suggest that mechanoregulation of emerin localisation affects the deposition of key regulatory histone marks and transcription during the VZ-CP transition, informing both the timing and outcome of neurogenesis (Figure 5G).

## DISCUSSION

Here, we identified LINC-mediated nuclear mechanotransduction as a key regulator of neuronal fate commitment in a complex model of human corticogenesis. We showed that, when insensitive to the mechanical forces associated with the VZ-CP transition, the timing of progenitor commitment is compromised, leading to premature VZ exclusion, differentiation, and biased neuronal fate acquisition. By combining a robust dox-dependent LINC perturbation approach and scRNA-seq analysis, we mechanistically linked this phenotype to altered functional signatures: ERK activity, density-sensing, and widespread epigenomic and transcriptomic disruption through mislocalisation of emerin at the NE.

The role of mechanosensation in informing tissue homeostasis has been characterised across a number of model systems during development (Guillot and Lecuit 2013). In such contexts, mechanically transduced signals typically drive cell extrusion and apoptosis, simultaneously maintaining tissue volume and canalising responses to extrinsic signals such as morphogenic gradients (Miroshnikova et al. 2018, Moreno et al. 2019, Akieda et al. 2019, Mcginn et al. 2021, Aoki et al. 2025). However, it remains unexplored whether such signalling mechanisms play analogous roles during the native establishment of cellular diversity. Recent work has shown that induction of apical constriction in CBO lumens leads to a bias towards horizontally oriented basal divisions, highlighting how progenitor crowding can drive changes in delamination dynamics (Marchenko et al. 2026). Strikingly, artificial induction of VZ crowding in this manner led to an enrichment in IP specification, consistent with the opposite phenotype observed in LINC-decoupled cells in our paradigm. While no effect on downstream neurogenesis was observed, only early neurogenic timepoints were assayed, without assessment of layer-specific fate specification. Our selective mosaic model facilitates full exploration of the downstream consequences of LINC-decoupling, allowing us to trace direct effects of mechanical disruption on the fate acquisition of affected cells.

Our observation that upper-layer neuron (ULN) specification was selectively impacted is consistent with work showing that the bulk of ULN-fated cells are the progeny of IPs (Tarabykin et al. 2001, Mihalas et al. 2016, Bury et al. 2024). However, this remains a point of contention in the field, particularly in the context of human neurogenesis (Lamonica et al. 2013, Vasistha et al. 2015, Coquand et al. 2022, Nowakowski et al. 2024, Hatanaka et al. 2024). In mice, where superficial neuronal layers are less reliant on the IP niche, studies leveraging double knockout of SYNE1/2 have reported an inversion of cortical layering, without loss of upper-layer neurogenesis (Zhang et al. 2010). In this *in vivo* study, the authors also observed premature apical progenitor depletion and a loss of basal mitosis consistent with a potential reduction in the IP population. However, the widespread consequences of complete SYNE knockout make direct comparison with our human organoid system difficult, as loss of regions beyond the CHD lead to additional impairment in microtubule motor recruitment and MT-directed migration (Zhu et al. 2017, Gonçalves et al. 2020).

Beyond the influence of cell-extrinsic tissue-scale forces, Nesprin-2-based interactions with all 3 classes of cytoskeletal components are a key pathway by which cell-intrinsic migratory forces are transduced to the nucleus (Grady et al. 2005, Yu et al. 2007, Zhang et al. 2007, Infante et al. 2018, Nakazawa & Kengaku 2020). While the SRK construct leaves the microtubule motor recruitment domain of the complex intact, it follows that excluding other endogenous nesprin isoforms from the ONM may provide an additional contribution to altered mechanotransduction. Dynein is a mediator of nucleo-centrosomal linkage, consistent with the premature exclusion from the VZ observed in SRK+ cells, as centrosomal recruitment to the apical plane of the tissue facilitates apical pulling forces during IKNM (Tsai et al. 2007, Tsai et al. 2010). Likewise, while neurons and basal progenitors are disseminated through the CP via a mix of migratory modes, many rely on nuclear translocation facilitated by centrosomal linkage, suggesting loss of physical connection to the NE at this stage contributes to the aberrant cortical positioning of SRK+ cells (Tanaka et al. 2004, Shu et al. 2004, Hansen et al. 2010, Wimmer et al. 2025). Interestingly, overexpression of similar nesprin constructs lacking the CHD has been shown to have minimal effects on migration and centrosome recruitment in mouse neurons, implicating LINC-dependent processes beyond centrosomal connection driving altered cortical stratification (Gonçalves et al. 2020).

The changes in ERK activity we observed in SRK-induced cells is consistent with previous work illustrating ERK’s importance during neurogenesis (Guillemot and Zimmer 2011, Kang and Herbert 2015). Specifically, ERK and upstream FGF-signalling have been shown to regulate the pace of RG to basal progenitor differentiation in mouse cortex (Kang et al. 2009, Sun et al. 2024). In humans, increased ERK activity drives generation of deep-layer neurons preferentially over more superficial layers, consistent with the inability of SRK-induced cells to produce upper-layer progeny (Gantner et al. 2021). Previous studies on neurogenesis have neglected ERK’s emerging role as a mechanotransducer, despite a range of literature showing altered ERK activity in response to mechanical cues spanning cortical tension, cell crowding and local tissue curvature (Aoki et al. 2013, Kawabata and Matsuda 2016, Hino et al. 2020, De Belly et al. 2021, Hino et al. 2022, Crozet and Levayer 2023, Hirashima and Matsuda 2024). Further literature has found direct interactions between nesprin-2 and ERK in the nucleus, suggesting recruitment of the factor to the nuclear envelope may influence pathway activity (Warren et al. 2010). Our findings show that ERK mechanoregulation plays a key role in modulating corticogenesis. Given the intersection of many of these activatory stimuli within the VZ, future experiments correlating ERK activation in LINC-decoupled cells with spatial tissue characteristics or precise mechanical measurements will be necessary to disaggregate the exact driver of ERK misregulation.

Our model suggests that the altered differentiation trajectory of LINC-decoupled cells is associated with cell-type specific changes in the deposition of the key histone variants H3K27me3 and H3K9me2,3. These data are consistent with work showing the importance of both marks in the temporal control of neurogenesis and maturation (Telley et al. 2019, Ciceri et al. 2024, Wu et al. 2022, Wu et al. 2025). Further literature has demonstrated these canonically repressive histone marks change their deposition and localisation in response to mechanical stimuli, both in the context of differentiation or mechanoprotective responses to acute stimuli (Demmerle et al. 2012, Le et al. 2016, Nava et al. 2020, Biedzinski et al. 2020, Miroshnikova and Wickström 2022, Hamouda et al. 2025). While multiple mechanisms have been proposed linking mechanical force to histone architecture, we observed relocalisation of the nuclear envelope protein emerin to the outer membrane, previously demonstrated in mouse epidermis and neuroectoderm (Le et al. 2016, Hamouda et al. 2025). These studies utilised disparate modes of mechanical perturbation, with the former applying biaxial cyclic stretch with a cell-stretcher device, and the latter relying on cell-intrinsic forces generated during the shape change accompanying priming (Bergert et al. 2021, De Belly et al. 2021). Our observation that LINC-decoupling leads to ONM enrichment of emerin, typically associated with increased nuclear strain, raises further questions about the exact nature of the forces acting on the nucleus during the VZ-CP transition.

While more representative of true tissue-scale mechanics, the forces explored in our experiments are a less controlled set of perturbations than those explored previously. The combination of cell-intrinsic and extrinsic mechanical signals, as well as variations in cytoarchitecture between or within VZ-CP transitions, compound to form the sum of mechanical stimuli experienced by any given nucleus in the tissue. In contrast with the unimodal stimuli probed in the studies referenced above, our data points to a global role for LINC-based mechanotransduction in informing bidirectional emerin switching in physiological contexts, rather than exclusively driving INM to ONM relocalisation. Given that our nuclear morphometry data shows the largest effect of SRK induction on nuclei occurs in the VZ, preceding the aberrant accumulation of H3K9me2,3 observed in the SOX2- population, it follows that premature loss of strain in the VZ lead may to aberrant compensatory ONM emerin switching upon crossing the VZ-CP boundary, analogous to processes of ‘mechanical memory’ shown in other systems (Beedle and Roca-Cusachs 2023, Scott et al. 2023, Kalukula et al. 2025, Lee et al. 2026). The lack of cytoplasmic markers compatible with digitonin permeabilisation precluded us from exploring whether this cell-type specific relocalisation was responsible for the observed effect on emerin in the NE.

In summary, our study demonstrates that mechanically transduced signals from the cortical tissue environment are a dynamic and fundamental component of developmental context. Through controlled disruption of a single mechanosensitive pathway in developing cells, allowed by our 3D organoid perturbation system, we gain unprecedented insight into the role of these signals in human cell fate decision-making. Our paradigm offers future possibilities to elucidate the precise role of mechanoregulation in neuronal differentiation and tissue patterning relevant to the human central nervous system.

### Limitations of this study

Given the morphological heterogeneity and limited spatial organisation in CBOs, we did not fully explore the impact of LINC decoupling on other cell types beyond those we examined. While, patterning of superficial cortical layers (I-III) and progenitor niches such as the outer or inner sub-ventricular zones may additionally be affected by SRK induction, our study provides insights into the principles of mechanotransduction-driven cortical neuronal cell type-specification.

Moreover, further experiments are required to confirm a direct link between the observed loss of IP fate and ULN specification in SRK+ cells. An interesting future avenue in this respect would be to investigate the effect of LINC-decoupling in mouse cortex, where the IP niche - and neuronal output thereof - is reduced and therefore a greater proportion of the ULN population is produced via direct neurogenesis from apical progenitors.

Finally, expressing the SRK construct under a single Tet-Responsive-Element array does not preclude the possibility of construct-silencing in some cells after induction. Therefore, the SRK- population may contain a small population of cells which expressed the construct prior to the 30-45DIV window, which could potentially mask the full effect of LINC-decoupling in the tissue.

## Resource availability

### Lead contact

Requests for further information and resources should be directed to and will be fulfilled by the lead contact, Alex Kingston.

### Materials availability

This study did not generate new unique reagents.

### Data and Resource availability

Upon publication, code for bioinformatic and image analyses will be available at Github. Raw sequencing data will be deposited at GEO under accession number GSEXXXXX. Any raw data or additional information required to re-analyse the data is available from the lead contact upon reasonable request.

### Declaration of generative AI and AI-assisted technologies in the manuscript preparation process

During the preparation of this work, the authors used Claude Sonnet for proofreading and optimising code written for bioinformatic and image analysis. The authors reviewed and edited the output as needed and take full responsibility for the content of the published article. No generative AI tools were used during writing of the manuscript.

## Acknowledgements

The authors thank Ben Steventon, Ruben Perez-Carrasco, Tony Southall, Oscar Baldwin, and all members of the Basu and Lakatos laboratories, for insightful discussion and advice during manuscript preparation. We thank Beata Orlos, Damien Box and Jo Jack for logistical and administrative support throughout the project. We thank Gabriela Grondys-Katarba and Reiner Schulte of the CIMR flow cytometry core for assistance with panel design. We thank all members of the CSCI imaging core, in particular Darran Clements, Louis Elfari and Olivia Hill, for their assistance with optimising imaging approaches. We thank all members of the CRUK sequencing core for sequencing assistance. We thank Matthew Williams, Hannah T. Stuart and Sumin Jang for assistance with developing fixed-cell cytometry protocols. We thank Luke Harland, Luke Bury, Yentel Mateo Otero and Emma Moth for advice with the scRNA-seq experimental design and analysis.

## Author contributions

Project leads: S.B. and A.L. Funding and conceptualisation: A.J.K., S.B., A.L. Experimental design, implementation, and analysis: A.J.K. Data interpretation: A.J.K., S.B., A.L. Technical help - scRNAseq library preparation: A.A.M.W. Cloning of the neGFP plasmid: A.S.K., K.M., A.J.K. Supervision: G.G.A., S.B., A.L. Writing – first draft: A.J.K., editing: S.B. and A.L. All authors commented on the manuscript.

## Funding information

A.J.K. and A.W. are funded by Wellcome Trust PhD studentships in Stem Cell Biology and Regenerative Medicine (218481/Z/19/Z). G.G.A. is funded by the UKRI Biotechnology and Biological Sciences Research Council (BB/W000423/1, BB/W000423/2). S.B. is funded partially by the Wellcome Trust (203151/Z/16/Z), UKRI Medical Research Council (MC_PC_17230), Trinity College Stem-Cell Medicine Senior Postdoctoral Researcher Fund and Imperial College London. A.L. and his team, including A.S.K. is supported by the UK Research and Innovation Medical Research Council (UKRI MRC Senior Clinical Fellowship; MR/X006867/1 awarded to A.L.).

## Competing interests

A.L. is a co-founder of Replicam Inc. All other authors declare no competing interests.

## METHODS EXPERIMENTAL MODEL

### Stem cell culture

Human Embryonic Stem Cells (hESCs; H9, WA09 WiCell and H9-SRK/rtTA-neGFP, generated as described below) were cultured in Stemflex (Thermo Fisher Scientific, A3349401) on Geltrex-coated 6-well plates (Thermo Fisher Scientific, A1413202) at 37°C and 5% CO_2_. Coating was carried out using Geltrex diluted 1:1000 in Dulbecco’s Modified Eagle Media (DMEM, 11966025). Cells were fed daily with 1.5-2mL prewarmed Stemflex and passaged 1:10-1:20 at ∼70% confluency (typically every 5 days). Colony passaging during regular culture was achieved using ReleSR (Stem Cell Technologies, 05872) according to manufacturers protocol. For applications requiring single-cell passaging, cells were dissociated using TrypLE (Thermofisher, 12604013) and reseeded at the desired density.

### Cortical organoid generation and culture

Human cerebral organoids with telencephalic identity were generated according to a modified version of the protocol previously described in Lancaster and Knoblich (2014), using the STEMdiff Cerebral Organoid Kit (StemCell Technologies, 08570). On day 0, hESCs at ∼70% confluency were treated with Accutase for 8-10 minutes at 37°C to generate a single cell suspension. The Accutase fraction was collected and transferred to a falcon tube containing 2ml DMEM F/12 (Thermofisher, 21331020). The suspension was spun down at 300g for 5 minutes, before resuspension in 1mL EB media supplemented with 10µM ROCKi (EB/Ri; Tocris 1254). Cells were counted and the volume of cell suspension required to obtain a cell density of 10-15000 cells per 100μl was then added to a fresh tube containing EB/Ri. 100μl of the resulting suspension was transferred to each well of an ultra-low-adhesion 96-well plate (Corning; CLS7007). On day 3 media was replaced with fresh EB medium without ROCKi, followed by neural induction (NI) media on day 5. On Day 7 individual organoids were moved to embedding sheets (StemCell Technologies, 08579) with a cut-tip pipette and embedded in 100% Matrigel (Corning CB-40234) before being transferred to 24-well ultra-low adherence tissue culture dishes (Corning; 353046) in Expansion medium. On day 10 Matrigel was removed by trituration and media replaced with Improved Differentiation Media with vitamin A (IDM+A; See table 1 for composition). Day 10 organoids were subsequently moved to a 60mm deep-walled tissue culture dish (Sigma-Aldrich; CLS430589) and placed on an orbital shaker (IKA; 0009019200) housed in an incubator for culture up to day 60. Media was exchanged every 3-4 days with fresh IDM+A. For induced KAS:H9 organoids, IDM+A was supplemented with doxycycline (dox) every 48h to a concentration of 3µg/mL from day 10 until the timepoint of interest. Chimeric organoids were generated by mixing cell suspension in ROCKi at a ratio of 70% SRK: 30% neGFP prior to EB seeding.

**Table 1:** Primary antibodies.

| Antigen | Species | Catalogue Number | Dilution IF | Dilution cytometry | RRID |
| --- | --- | --- | --- | --- | --- |
| MAP2 | Chicken | ab5392 (Abcam) | 1:1000 | — | AB_2138153 |
| SATB1/B2 | Mouse | ab51502 (Abcam) | 1:200 | — | AB_882455 |
| CTIP2 | Rat | ab18465 (Abcam) | 1:500 | — | AB_2064130 |
| SOX2 | Goat | AF2018 (R&D Systems) | 1:500 | 1:2000 | AB_355110 |
| Vimentin | Rabbit | ab92547 (Abcam) | 1:200 | — | AB_10562134 |
| Eomes (TBR2) | Rabbit | ab216870 (Abcam) | 1:500 | — | AB_2943124 |
| TBR1 | Rabbit | ab183032 (Abcam) | 1:200 | — | AB_2936859 |
| RFP | Rabbit | ab124754 (Abcam) | 1:500 | 1:1000 | AB_10971665 |
| RFP | Rat | 5f8 (Proteintech) | 1:200 | 1:500 | AB_2336064 |
| H3K9me3 | Mouse | 5327 (Cell Signaling) | 1:200 | 1:2000 | AB_10695295 |
| H3K27me3 | Rabbit | 9733 (Cell Signaling) | 1:200 | 1:2000 | AB_2616029 |
| TPX2 | Rabbit | NB500-179 (Novus) | 1:500 | — | AB_10002747 |
| Nestin | Mouse | ab22035 (abcam) | 1:500 | — | AB_446723 |
| Phospho-ERK1/2 (Thr202/Tyr204) | Rabbit | 9101 (Cell Signaling) | — | 1:3000 | AB_331646 |
| RNAPII-S2 | Rabbit | ab5095 (Abcam) | — | 1:1000 | AB_304749 |
| Emerin | Mouse | H-12 (Santa Cruz Biotechnology) | 1:100 | 1:100 | AB_2100051 |
| Lamin B | Rabbit | ab229025 (Abcam) | 1:500 | 1:500 | AB_3083735 |

### Plasmids and Generation of KAS:H9 line

The control neGFP plasmid sequence was obtained from a TRE:SUN1-TorsinNELS-eGFP template, which contains an identical backbone to the SRK plasmid (Luxton et al. 2010). Fragments were cloned using the In-Fusion cloning kit (638948, Takara Biosciences) to remove the SUN1 sequence, resulting in a piggybac-compatible backbone containing the Tet-Response-Element array and neGFP protein sequence. The SRK and rtTA:neGFP cell line were generated by two-step transfection of H9 cells with the SR-KASH/neGFP plasmids and an rtTA doxycycline response element (SRK and rtTA plasmids kindly supplied by Kevin Chalut). Transfections were carried out using the Lipofectamine STEM reagent kit (Invitrogen; STEM00003) according to an adapted version of the kit protocol. Briefly, 24 hours prior to transfection, low passage number H9 cells were single-cell passaged onto 24-well plates as described above at a density of 50000 cells/well. The following day, the culture media was replaced with OPTI-MEM (Gibco; 31985-062) immediately prior to incubation with liposomes containing the desired plasmid and an equal amount of the PiggyBac transposase plasmid to facilitate transposition into the host genome. Cells were incubated in the liposome suspension for 4 hours before dilution with 500μl Stemflex, the media was subsequently exchanged for fresh stemflex after 12 hours. For each experiment one control well was prepared by incubation with liposomes without plasmid DNA. After 24 hours, the transfected colonies were passaged onto Geltrex-coated 10cm dishes (Thermofisher; 150350) for antibiotic selection. Colonies were allowed to grow for 24 hours before addition of selection antibiotics to the culture media (Hygromycin 10μg/ml, Blasticidin 2μg/ml), media was refreshed daily. Colonies with healthy hESC morphology remaining on the transfected plate were picked after all colonies on the control plate had died. Selected colonies were passaged onto individual wells of a 24-well plate for subsequent expansion and banking. 4 KAS:H9 clones were picked and subsequently validated for pluripotency and SRK expression via morphological analysis, FACS and Immunofluorescence-based marker expression. 12 neGFP clones were picked and selection carried out based on GFP expression and competency to generate CBOs.

## METHOD DETAILS

### Organoid cell dissociation

Organoids at 30 and 45 DIV were transferred to 10cm dishes containing Hank’s Balanced Salt Solution without calcium or magnesium ions (HBSS -/-; Gibco 12082739) and washed 2-3x. At this step, d45 organoids were bisected into smaller pieces using a sterile scalpel, to aid the removal of necrotic cells from the core of the tissue. After washing, organoid pieces were transferred to C-tubes (Miltenyi, 130-093-237) containing 2mL pre-equilibrated Papain reconstituted in HBSS +/+ (Worthington, LK003178; 20 units/mL) and ran on a gentleMACs Octo Dissociator (Miltenyi, 130-095-937) set to the ‘ABDK 30’ standard program at 37°C. After dissociation, the organoid suspension was gently triturated 10-20x using a p1000 to break up residual clumps of tissue, before addition to 2mL 1mg/mL DNAse solution (Sigma, 11284932001; reconstituted in PBS +/+ and activated for 5’ at 37°C) in a 15mL falcon tube. This suspension was spun down at 350g for 5 minutes and gently resuspended in another 1mL DNAse solution, before dilution with 5mL PBS -/-. The spin was repeated and, after resuspension in PBS -/-, the suspension was passed through a 30µm filter into a fresh tube. A final spin at 450g for 5 minutes was used to pellet the filtrate. The following steps of the protocol depended on the intended use of the cell suspension. For scRNAseq, the pellet was resuspended in 1mL PBS -/- +0.04% BSA (Bovine Serum Albumin; cat) and counted before proceeding to downstream library preparation or fixation. For intracellular cytometry, resuspended cells in PBS -/- +0.5% BSA were counted and split equally into 5mL polystyrene FACS tubes (cat) in 50µl aliquots (targeting 500,000-5M cells/tube) prior to PFA or methanol fixation. For 2D culture, cells were resuspended in N2B27 + 1:1000 ROCKi before seeding on coverslips at the desired density.

### Immunohistochemistry

For histological processing, organoids were fixed in excess 4% paraformaldehyde (PFA; Sigma-Aldrich; 158127) overnight at 4°C on a tube roller or for 60 minutes at room temperature, then washed in PBS. Samples for cryostat sectioning were incubated for 24-72h in 30% sucrose in PBS at 4°C before embedding in Optimal Cutting Temperature compound (OCT; Thermofisher; 15212776). OCT blocks were subsequently cryosectioned at 10-12μm thickness on a Leica CryoJane (cat) and collected on superfrost slides (Thermofisher, 10149870). Sections were washed in PBS (3x10 mins) before blocking and permeabilisation in 0.3x Triton-X (PBS-T; Sigma-Aldrich, x100) in PBS with 10% serum (donkey or goat) for 1 hour. Primary antibodies were then applied overnight at 4°C in 0.1x PBS-T with 3% serum. The following day slides were once again washed with PBS (3x10 minutes) before secondary antibody incubation in PBS + 1µg/mL DAPI (Sigma-Aldrich; D9542-1MG) for 1 hour at RT in the dark. Finally, slides were washed for 3x10 minutes in PBS. Sections were mounted in FluorSave (Sigma-Aldrich; 345789) or Prolong diamond (ThermoFisher; P36965) and stored at 4°C until imaging. Details of primary and secondary antibodies used for staining can be found in tables 2 and 3.

**Table 2:** Secondary antibodies.

| Antigen | Dilution IF | Dilution cytometry | Catalogue Number | RRID |
| --- | --- | --- | --- | --- |
| Donkey anti-Goat 488 | 1:500 | 1:3000 | ab150129 (Abcam) | AB_2687506 |
| Donkey anti-Rabbit 568 | 1:500 | 1:3000 | ab175692 (Abcam) | AB_3076303 |
| Donkey anti-Rat 488 | 1:500 | 1:3000 | A21208 (Invitrogen) | A-21208 |
| Donkey anti-Goat 647 | 1:500 | 1:3000 | A32849 (Invitrogen) | AB_2866498 |
| Donkey anti-Chicken 647 | 1:500 | 1:3000 | A78952 (Invitrogen) | AB_2921074 |
| Goat anti-Rabbit 647 | 1:500 | 1:3000 | ab150075 (Abcam) | AB_2752244 |
| Phalloidin 488 | 1:400 | - | A12379 (Invitrogen) | - |
| Phalloidin 647 | 1:400 | - | A22287 (Invitrogen) | - |
| Donkey anti-Rabbit 647 | 1:500 | 1:3000 | ab150075 (Abcam) | AB_2752244 |
| Donkey anti-Rat 568 | 1:500 | 1:3000 | ab175475 (Abcam) | AB_2636887 |
| Donkey anti-Rat 647 | 1:500 | 1:3000 | ab150155 (Abcam) | AB_2813835 |
| Donkey anti-Mouse 568 | 1:500 | 1:3000 | ab175472 (Abcam) | AB_2636996 |
| Donkey anti-Mouse 647 | 1:500 | 1:3000 | ab150107 (Abcam) | AB_2890037 |
| Donkey anti-Mouse 750 | — | 1:3000 | ab175739 (Abcam) | AB_2936438 |

### Flow cytometry

For flow cytometry based on intracellular antigen markers, cell suspensions were prepared as described above, with the exception of using Accutase as a dissociation reagent, and simple manual trituration without use of an automated dissociator. Cells were fixed with 4% PFA for 15 minutes on ice. Fixed cells were spun down at 1000g for 5 minutes and resuspended in PBS -/-. Cells were spun again and the supernatant aspirated until 50-100µl remained, the pellet was resuspended in this volume before dropwise addition of 100% methanol pre-chilled to -20°C while agitating cells on a vortex at low speed to avoid clumping. Methanol permeabilised cells were stored at -20°C until use. For staining, cells were spun down and resuspended in ice-cold PBS+0.5% BSA. The spin down and resuspension was repeated a further 2 times for a total of 3 washes. Cells were then resuspended in 100µl of primary antibody diluted in PBS+0.5% BSA and incubated overnight at 4°C on the orbital shaker. Cells were then spun down and washed 3 times as detailed above, before resuspension in 200µl secondary antibody in PBS+0.5% BSA + 1:3000 DAPI and incubation for 30 minutes in the dark. Cells were then spun down and washed for a final 3 times before resuspension in PBS+0.5% BSA for cytometry. Unstained, single stain and no primary control tubes were included in all replicates for all panels. For emerin localisation experiments, the methanol addition step was omitted to prevent non-selective membrane permeabilisation. Cells were therefore stored in PBS+0.5% BSA at 4°C prior to staining. Staining was carried out using the above protocol for both fixation methods. Details of primary and secondary antibodies used for staining can be found in tables 2 and 3. For quantifying the expression of intracellular markers, a BD LSRFortessa was used to quantify expression of intracellular markers. Five lasers (355nm, 405nm, 488nm, 561nm, 640nm) and a 100µm nozzle were used to acquire flow cytometry data from PFA-fixed, methanol-permeabilised samples dissociated as described above. A minimum of 10,000 events were collected per sample, after gating for debris and doublets based on FSC-A/SSC-A, FSC-A/FSC-H and DAPI. No-primary controls were also included for each panel to set gates for positive signal. To prevent surviving signal from endogenous RFP interfering with other channels, the 561 channel was reserved for ant-RFP antibody staining in all panels. Compensation, gating and downstream data analysis were performed in the FlowJo software package (v10/v11). Samples with fewer than 1000 SRK+ cells after gating were discarded.

### Fluorescence-activated cell sorting (FACS)

After dissociation to single cell suspensions as described above, organoid samples were resuspended to the density of 2-15M cells/mL in cold PBS+0.5% BSA. A BD FACSAria Fusion cell sorter was used to SRK+ and SRK- cells based on expression of RFP. Uninduced control samples were used to establish gating for debris and doublets based on FSC-A/SSC-A and FSC-A/FSC-H, as well as the SRK- population gate. Induced (doxycycline-treated) samples were then run through the instrument and populations segregated based on RFP signal (561nm laser), with all events exhibiting RFP emission higher than the previously established SRK- gate retained as SRK+.

### Image acquisition

Brightfield images of CBOs were acquired on a Leica DMi8 inverted microscope (10x, 5x objectives) or a Thermofisher EVOS m3000 (2.5x, 4x, 20x objectives; Brightfield and RFP filter cubes). Confocal images of nuclear morphology, marker expression and SRK/neGFP distribution were acquired using a Zeiss 980 microscope, using 20x, 40x and 63x objectives. Acquisition details for each experiment are described below. Where appropriate, consistent laser power and acquisition settings were used for each experiment.

### Single-cell RNA-seq library preparation

Libraries for single-cell RNA sequencing of KAS:H9 organoid development were prepared using the Parse Biosciences cell fixation and Evercode whole-transcriptome v3 kit (ECFC3300, ECWT3300), according to the manufacturer’s protocol. Sublibraries were pooled at equimolar concentrations and sequenced on a single lane of an Illumina Novaseq X to a target depth of 60,000 reads/cell (3.1Bn reads).

### Single-cell RNA-seq analysis

Raw FASTQ files generated from illumina sequencing were processed using the Trailmaker pipeline module (v1.5.0, parse biosciences). Sample-specific barcodes were demultiplexed using a custom script to allow separation of human and mouse samples before mapping to the respective custom reference genomes. Custom reference genomes (hg38) were generated by appending the bottom-strand sequence of the original mRFP sequence from the SRK plasmid to the FASTA file of the corresponding genome. A matching GTF annotation was then added to the appropriate genome index file. Reference genome mapping was performed in Trailmaker, and the filtered count matrices subsequently analysed in Seurat (v5). Background reads were removed by setting a minimum transcripts per cell threshold on a per sample basis based on the point of inflection in a knee plot (threshold range: 1227 to 2550 transcripts/cell). Dead or dying cells were removed by filtering barcodes with high mitochondrial content (3%). Outliers in the distribution of number of genes vs number of transcripts were removed by fitting a spline regression model. Additionally, any cell expressing less than 500 or more than 17500 features was excluded from further processing. Cells with a high probability of being doublets were filtered out using the scDblFinder method using standard parameters (Germain et al. 2022). For normalisation and clustering, the data were log normalised and the 1500 most highly variable genes selected (selection method ‘vst’) for scaling and principal component analysis. Scaling was carried out on each timepoint individually to remove technical bias. Neighbour identification and clustering (Leiden algorithm) was performed on the top 30 PCs, selected based on visual inspection of an elbow plot. A cluster resolution of .5 was selected for preliminary analysis. Integration across timepoints was then carried out using the Seurat v5 implementations of canonical correlation analysis (CCA), harmony integration (Korsunsky et al. 2019) and reciprocal PCA integration (rPCA). UMAP embeddings from each integration were reclustered and benchmarked using the SeuratIntegrate package, from which CCA was identified as best preserving biological structure in the data. Stressed cells were filtered out of the dataset using the Gruffi R package, before renormalisation and reclustering. A single cluster of non-telencephalic cells (choroid plexus) was identified after cluster annotation by expression of the canonical markers TTR, LMX1A, OTX2 and PRLR. This cluster was removed from downstream analysis to better preserve relevant forebrain differentiation trajectories. CCA clusters were annotated using a combination of canonical marker expression and projection onto a reference atlas of fetal human brain development (Qian et al. 2025). The former was accomplished using Seurat’s FindAllMarkers function, with a minimum percentage threshold of .25 and log fold change threshold of .5. Integration and Label transfer was accomplished as described below (mosaic integration and spatial imputation). Ultimately, projection and manual marker assessment were assessed in parallel to assign final cell type annotations to CCA clusters. For SRK- expressing cell identification, cells in the dox treated condition expressing 1 or more SRK-mapping reads were assigned as SRK+. SRK-expressing cells in the control condition were excluded from further analysis, after immunofluorescent assessment of organoids from the batch used for sequencing confirmed an absence of any SRK protein expression in cortical organoids. Cluster-specific enrichment of SRK+ cells was assessed using Fisher’s exact test against the proportion in the whole dox-treated sample. Pseudotime and differential abundance analysis were performed using the monocle3 and milo packages, respectively (Trapnell et al. 2014, Dann et al. 2022). Differential abundance was assessed across condition and timepoint, with both models including batch as a covariate during matrix design. For transcription factor scoring, ERK and YAP targets were curated from a combination of published literature and KEGG/GO databases (GO:0070371, GO:0035329, Cordenonsi et al. 2011, Chen et al. 2023). Genes were filtered for expression (at least 50% expression in any single cluster). The resulting genes were used to generate a compound score with the seurat AddModuleScore() function. Scores were aggregated per sample and modelled by linear regression including timepoint, SRK status, and their interaction (score ∼ timepoint × SRK status); p-value reflects the timepoint:SRK interaction term. The summary of genes for each score were included in Table 4.

### Mosaic integration and spatial imputation

For integration into the merFiSH atlas, we leveraged a mosaic approach, optimised for integration between datasets with a large proportion of unshared features (Ghazanfar et al. 2021, Harland et al. 2024). The shared high-dimensional embedding produced by mosaic integration was then batch-corrected with harmony and resulting matrix used to generate UMAP embeddings combining both datasets. For label transfer of atlas labels to the organoid data, we utilised an adaptive K-nearest neighbours approach, with values of k between 5-55 tested for each cell. For continuous variables such as relative cortical height a similar approach was used, but with the mean of k neighbours assigned to the query cell. Local density calculations were carried out on the merFiSH spatial embeddings, with local density defined as the inverse of the mean distance to the 10 nearest cell centroids in euclidean space. This value was then used to impute the local density of organoid cells with the same approach described above. Visualisation of organoid cells on fetal brain spatial data was achieved simply by transfer of spatial coordinates from the nearest merFiSH cell in each sample to the query cell in question, these values were used for visualisation only, with all quantification of spatial position utilising properly imputed values. The tradeseq package was used to extract density-responsive genes by fitting a generalised additive model to gene expression of highly variable features against imputed local density. The associationTest() function was then used to identify genes which varied significantly with density, using an FDR-adjusted p-value of .05 as a threshold. Genes were then grouped into clusters using the ClusterExperiment package with a minSize parameter of 50, which identified 11 total expression patterns. The 4 largest response clusters were selected for further analysis, see supplementary table 1 for a full list of genes in each cluster.

### Image processing and analysis

For nuclear morphometrics, 63x Z-stacks of VZ-CP boundaries were acquired at 0.4µm axial resolution, spanning the entire region of DAPI signal to prevent truncation of cells in the Z-lane. The DAPI channel was subsequently segmented using Cellpose SAM (v4.0.1; Pachitariou et al. 2025), with all 2D orthoplanes segmented per stack. 2D segmentation outputs in all orthogonal planes were then stitched into cohesive 3D objects using uSegment-3D (Zhou et al. 2025). Poor quality masks and objects touching the XY boundary were discarded prior to downstream analysis. For immunofluorescence and confocal microscopy image acquisition, exposure parameters were kept the same. For quantitative analyses, unmodified images were used and processed in FiJI (v2.16.0). For illustration purposes, the recommended guidelines were followed. Images were uniformly and only minimally processed by adjusting exposure parameters within and across images in Fiji without affecting data presentation. Morphometrics were calculated using a custom script in Rstudio, prior to manual VZ-CP boundary annotation based on SOX2 signal, and identification of SRK+ cells using Gaussian-Mixed-Model thresholding. Distance calculations were based on the minimum euclidean distance of the nuclear centroid to the tissue boundary. For assessing VZ exclusion and SRK+ cell position, 20x images of full VZs were acquired at timepoints spanning 30-45DIV. All SRK or neGFP+ cells per ROI were identified and relative VZ position calculated using a custom python script. Only VZs containing both SRK and control cells were selected for analysis. For overlap with TBR2+ IPs non-chimeric organoids were assessed and costained for TBR2 at 30 and 45DIV. To evaluate IP and neuronal marker coexpression, 20x images of full VZs were acquired at 15, 30, 45 and 60DIV. 3x 100x100µm ROIs per image were randomly selected and cell-type proportions manually counted within each ROI. Results were pooled and averaged per image and proportion of positive cells per population calculated. Apical RG division angle was quantified using 40x images of TPX2 and nestin immunostaining of the mitotic spindle and apical VZ boundary, respectively. The angle tool in FiJI was used to measure the acute angle between the VZ apical surface and the mitotic axis. Cells where one centrosomal pole was clearly visible were measured as perpendicular, with the corresponding pole assumed to be orthogonal to the image plane.

## STATISTICAL ANALYSIS AND REPRODUCIBILITY

### Statistical details

Statistical analysis was performed in RStudio. A summary of statistical approaches and sample sizes can be found in supplementary table 2. Where appropriate, a Shapiro-Wilk test was used to assess normality of observation distributions, non-parametric tests were applied when the null was rejected. Throughout all experiments, at least 6 organoids spanning 3 independent differentiations were used for statistical analysis, with observations pooled by organoid for comparisons, with the exception of the chimeric organoid analysis in Figure 2d, where statistics compare individual VZs.

### Supplemental information

**Supplementary Table 1:** Gene score lists, primer sequences used for Infusion cloning

**Supplementary Table 2:** Statistical details and sample sizes

