## Supplementary figures 1-4 for "Organoids reveal niche-specific mechanotransduction-guided human cortical patterning and cell fate acquisition"

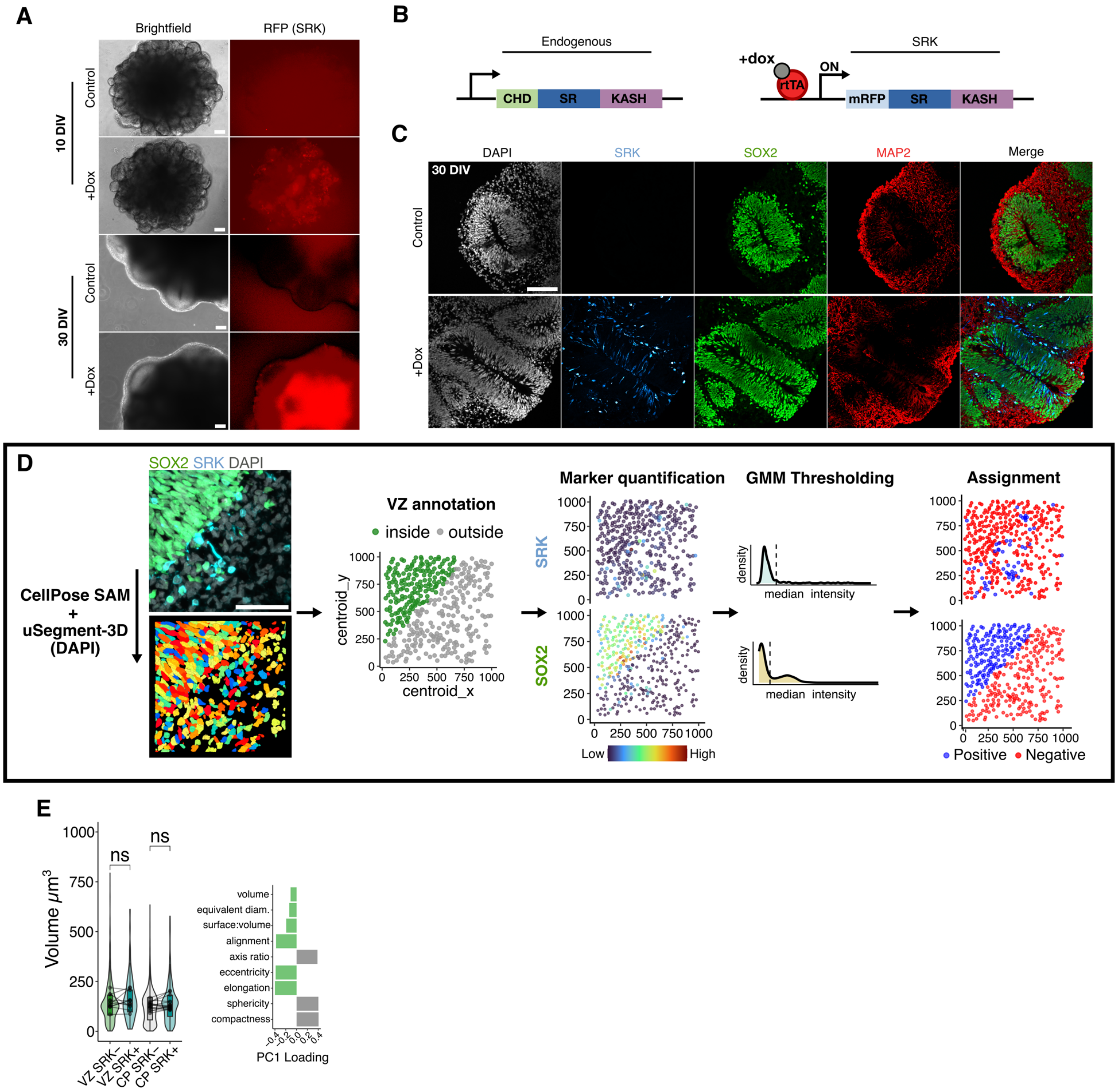


**Supplementary figure 1**: **Niche-specific nuclear morphology is disrupted by LINC-decoupling.**

(A) Brightfield (left) and widefield fluorescence images of induced and uninduced organoids during development, showing SRK expression through RFP signal. Scale bars 100µm.

(B) Schematic of endogenous and SRK constructs showing protein domains. CHD = calponin homology domain; SR = spectrin repeats, KASH = Klarsicht, ANC-1, spectrin-homology; mRFP = monomeric red fluorescent protein; rtTA = reverse tetracycline-controlled transactivator; Dox = doxycycline.

(C) Representative immunofluorescence (IF) images showing uninduced and dox-treated organoids at 30DIV. Scale bar 100µm.

(D) Schematic of nuclear morphometry analysis pipeline. GMM = gaussian mixed model; Scale bar 100µm.

(E) Violin plots showing nuclear volume across tissue compartments in SRK+ and SRK- cells (left) and top morphometric contributors to PC1 (right). *n* = 6 organoids (12 tissue boundaries from 3 independent differentiations). Paired two-tailed student's t-test. *p* = 0.25 (VZ) 0.38 (CP)

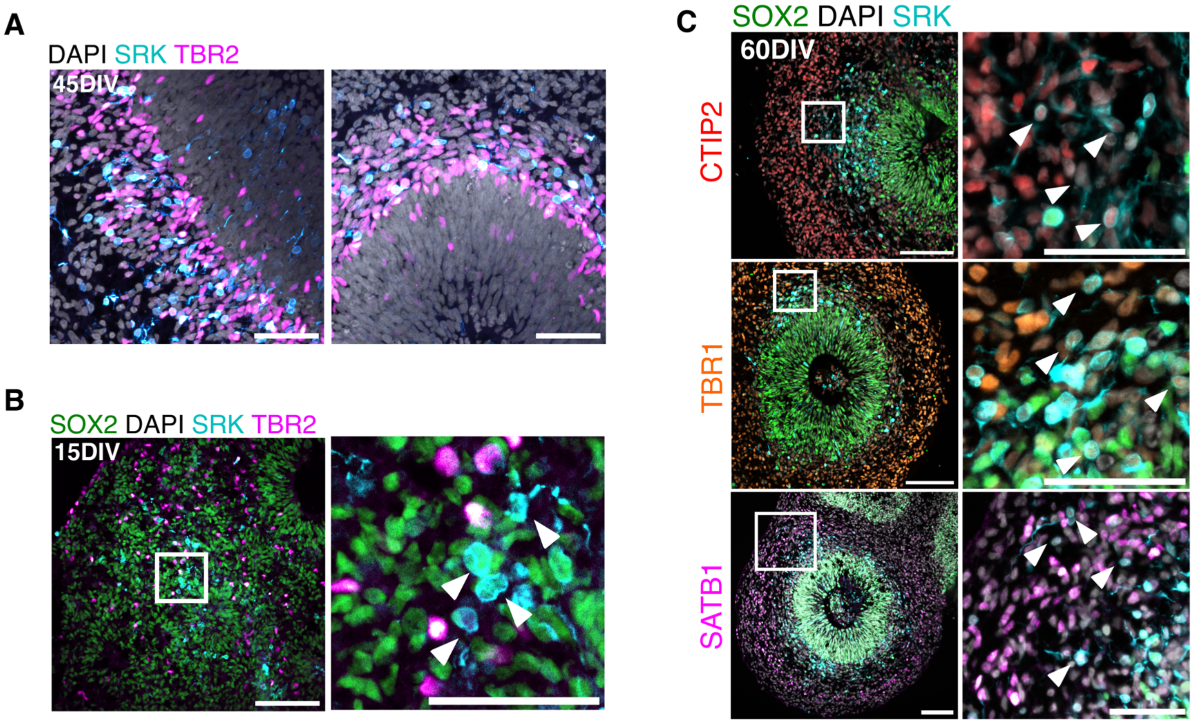


**Supplementary figure 2: LINC-decoupled cells are not competent to produce IPs and display biased neurogenic potential**

(A) Representative confocal images at 45DIV showing SRK and TBR2 expression in the organoid SVZ. Scale bars 100µm. Images representative of *n* = 3 independent experiments.

(B) Representative confocal images at 15DIV showing loss of TBR2 positive SRK-expressing cells at earlier timepoints. Scale bars 100µm (left) 50µm (right). Images representative of *n* = 3 independent experiments.

(C) Representative images of TBR1, CTIP2 and SATB1 expression, co-stained for SRK. Scale bars 100µm (overview) 50µm (inset).

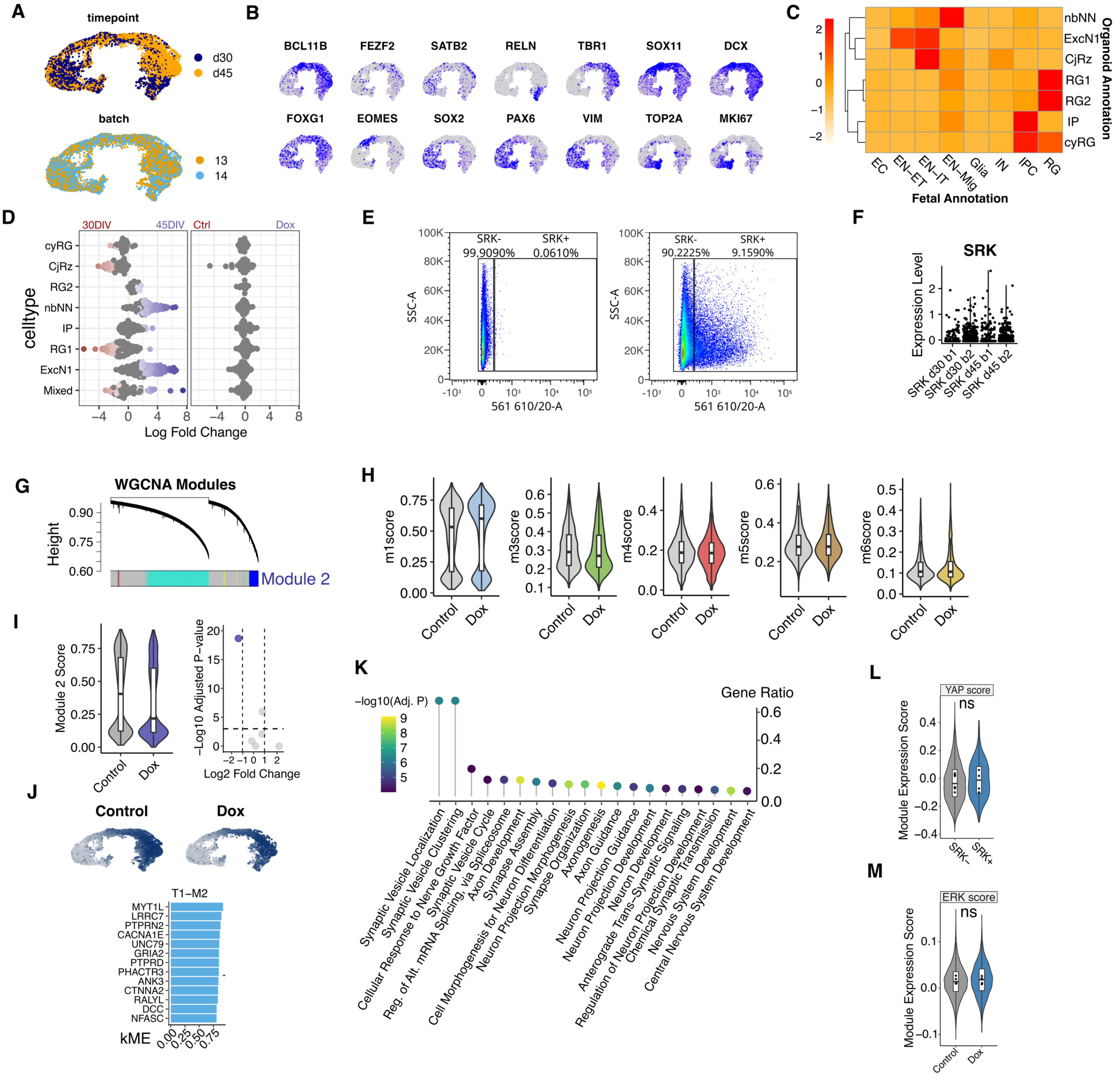


**Supplementary Figure 3: scRNAseq reveals altered differentiation trajectories in LINC-decoupled cells**

(A) CCA-integrated UMAP embedding of organoid scRNAseq data coloured by DIV and batch.

(B) UMAPs coloured by expression of key cell type markers. BCL11B, TBR1, FEZF2 (DLNs); SATB2 (ULNs); RELN (Cajal-Retzius cells); SOX11, DCX (Newborn neurons), EOMES/TBR2 (IPs); SOX2, PAX6, VIM (Radial Glia); TOP2A, MKI67 (cycling progenitors) FOXG1 (forebrain identity) .

(C) Heatmap showing mapping scores of organoid cell types (y-axis) to corresponding fetal cell types from Qian et al. (2025) pre-frontal cortex samples.

(D) Milo beeswarm plots showing neighbourhood differential abundance of cells from each timepoint (left) and treatment (right) in each cluster. Coloured points show significant enrichment *p_adj* < 0.05. General linear model with spatial FDR correction.

(E) Representative FACS plot of control (left) and induced (right) samples, showing degree of mosaicism in induced samples consistent with SRK detection in scRNA-seq data.
(F) Normalised expression of SRK transcripts per dox-treated sample from scRNA-seq data.
(G) WGCNA dendrogram showing all identified modules after co-expression analysis.
(H) Violin plots showing WGCNA module expression (excluding module 2)between treatment conditions.
(I) Violin plot (left) and volcano plot quantifying downregulation of module 2 in dox-treated organoids relative to uninduced samples.
(J) UMAP embeddings showing composite module 2 expression per cell in each condition (top) and top eigengene contributors (bottom)
(K) Lollipop plot of ENRICHR GO term analysis, showing involvement of module 2 component genes in neuronal maturation processes.
(L) Violin plot comparing ERK score between treatment conditions. $N = 4$ samples from 2 independent differentiations. Linear model with age as covariate, $p = 0.95$.
(M) Violin plot comparing YAP score between SRK+ and SRK - populations. $N = 4$ samples from 2 independent differentiations. Linear model with age as covariate, $p = 0.38$.
*p*-values: * <.05, **<.01, ***<.001; Overlaid box plots show median values (line), interquartile range (box) and data range excluding outliers (whiskers).

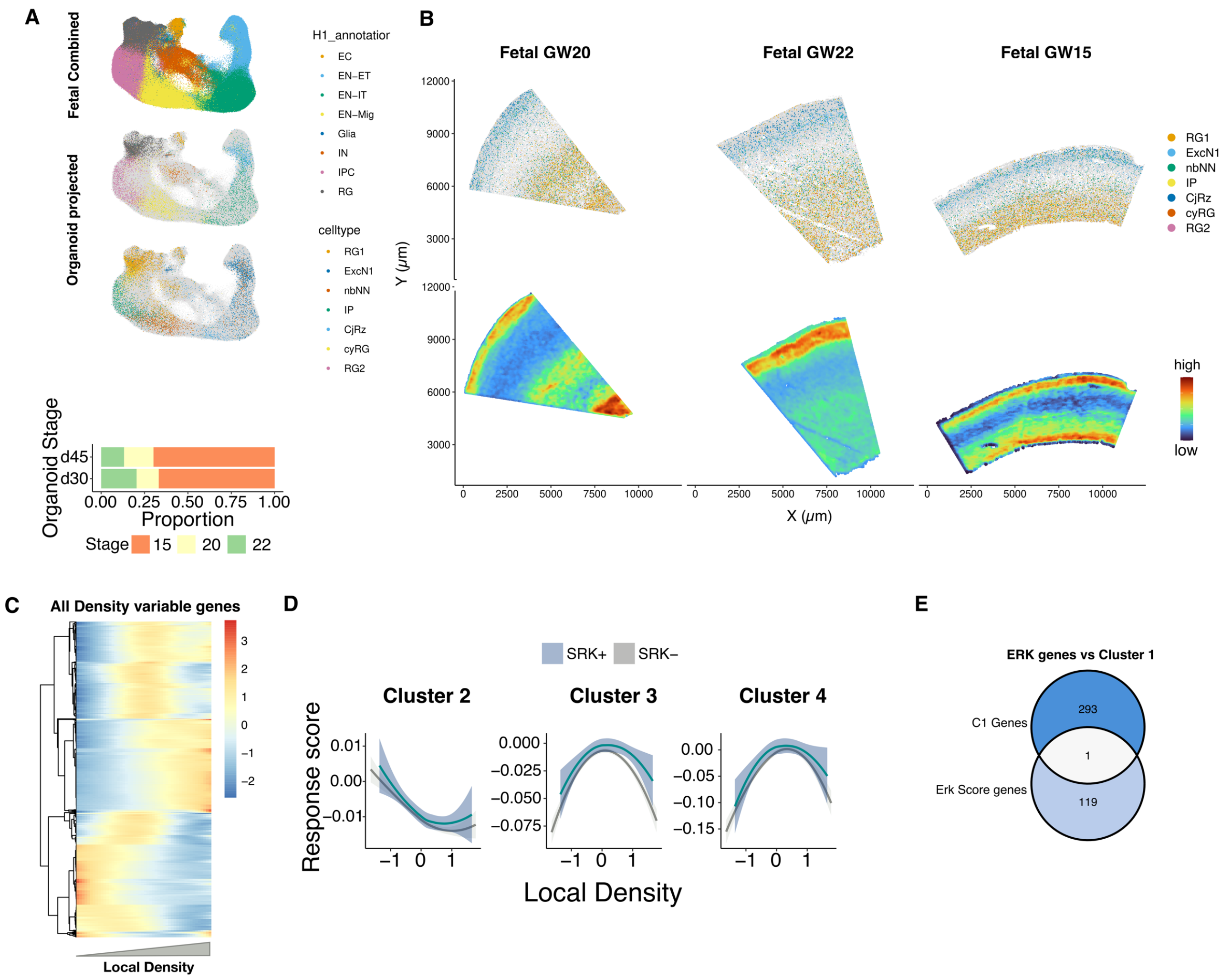


**Supplementary Figure 4: scRNAseq reveals loss of ERK-mediated crowding response in LINC-decoupled cells**

(A) Harmony-integrated UMAP projection of merFISH data coloured by fetal atlas H1 annotation (top). Organoid cells (coloured) projected onto merFISH UMAP, coloured by predicted fetal annotation (middle-upper) and organoid cell type label (middle-lower). Stacked barchart depicting percentage predicted fetal timepoint by organoid sample DIV (bottom).

(B) Spatial embeddings of all merFISH samples used for reference mapping, showing projected organoid cells coloured by cell type (top) and heatmap showing local density score across the tissue section (bottom).

(C) Heatmap showing all genes varying significantly with local density, as identified by tradeSeq, coloured by normalised expression score.

(D) Response scores of clusters 2-4 in SRK+ and SRK- cells in dox organoids plotted against local density. Ribbons show SEM, loess fit.

(E) Venn diagram depicting overlap of response cluster 1 with ERK score genes. Only 1 shared gene (<1%) was identified.

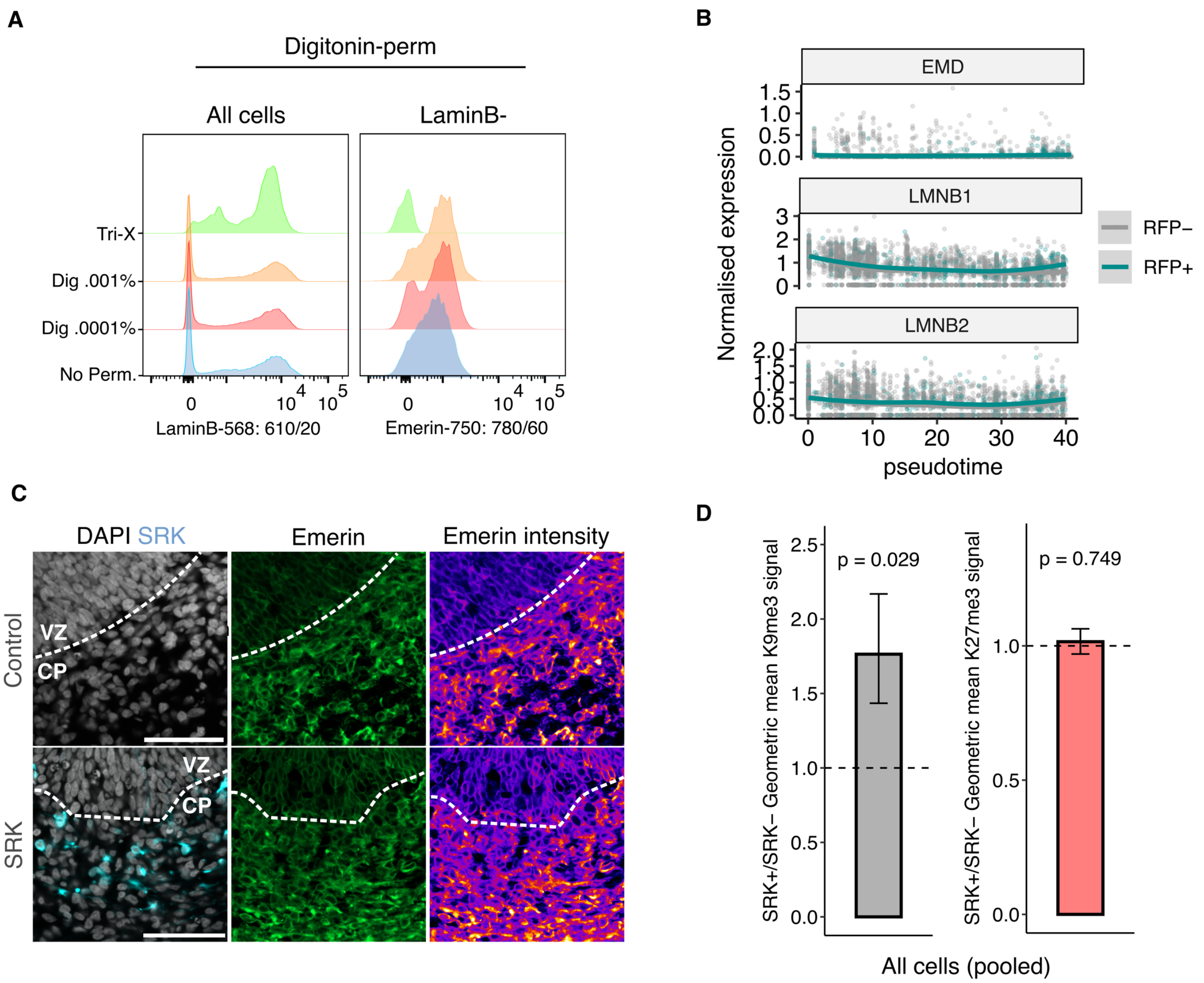


**Supplementary Figure 5: LINC decoupling drives widespread epigenomic dysregulation via Emerin mislocalisation**

(A) Confocal images of emerin expression across the VZ-CP boundary in induced and uninduced tissues, showing increased expression in the CP in both treatments. Scale bar 100µm.

(B) Scatter plots showing normalised expression of canonical NE proteins emerin and B-type lamins over pseudotime in SRK+/SRK- cells.

(C) Ridgeplots showing LaminB signal in all cells (left) and emerin signal in non-permeabilised cells (right) across all tested permeabilisation conditions, showing emergence of LaminB- population and retention of emerin signal at 0.001% digitonin treatment. Representative plot from N = 3 independent experiments.

(D) Barplots showing mean fold-enrichment of H3K9me2,3 (left) and H3K27me3 (right) IR in all SRK+ cells vs SRK- cells. Samples from $n$ = 4 independent differentiations. Unpaired two-tailed student's t-test. $p$ = 0.029 (H3K9me2,3), 0.749 (H3K27me3). Whiskers show SEM).
